# Role of the SUT1 and SUT2 sugar transporters during stolbur phytoplasma infection in tomato

**DOI:** 10.1101/2020.09.25.309708

**Authors:** Federica De Marco, Brigitte Batailler, Michael R. Thorpe, Frédérique Razan, Rozenn Le Hir, Françoise Vilaine, Alain Bouchereau, Marie-Laure Martin-Magniette, Sandrine Eveillard, Sylvie Dinant

## Abstract

Phytoplasmas inhabit phloem sieve elements and cause abnormal growth and altered sugar partitioning. But how they interact with phloem functions is not clearly known. The phloem responses were investigated in tomato infected by ‘*Candidatus* Phytoplasma solani’, at the beginning of the symptomatic stage of infection, both in symptomatic and asymptomatic leaves, the first symptoms appearing in the sink top leaf at the stem apex. Antisense lines impaired in the phloem sucrose transporters SUT1 and SUT2 were included. The infection in source leaves was not associated with symptoms. In the symptomatic, sink leaf, yellowing and leaf curling was associated with higher starch accumulation and expression of defense genes. The transcriptional analysis of symptomatic leaf midribs indicated that transcript levels for genes acting in the glycolysis and peroxisome metabolism in infected plants differed from these in non-infected plants. Phytoplasma multiplied actively in at least three additional lower leaves although they were symptomless, with no sign of activation of defense markers, although the rate of exudation of sucrose from these symptomless, source leaves was lower for infected plants. A few metabolites in phloem-enriched exudate were affected by the infection, such as glycolate and aspartate, and some of them were also affected in the control *SUT1*- and *SUT2*- antisense lines, in which sucrose retrieval or release in the sieve elements are impaired. A metabolic switch could explain the delivery of more glycolate into the sieve elements of infected plants. The findings suggest a link between sugar transport and redox homeostasis.

**One sentence summary:** An impairment of sucrose retrieval and release in the sieve elements occurs during phytoplasma infection, associated with changes in sugar and peroxisome metabolism

## Introduction

In their host plants, phytoplasmas represent an interesting case of obligate bacterial pathogens inhabiting the sieve elements (SE), which are transmitted by phloem-feeding insects. Phytoplasmas cause diseases affecting crops worldwide, provoking huge economic losses (Hollingsworth et al., 2008; Bertaccini et al., 2014). Because they multiply exclusively in the SEs, phytoplasma propagate systemically from the site of infection to sink organs, a process largely explained by convection along with the assimilate flow in the phloem (Siddique et al., 1998; Wei et al., 2004), even if the movement of phytoplasmas cannot be solely explained by mass flow (Christensen et al., 2005; Pagliari et al., 2017). The infection dynamic depends on the plant-phytoplasma pathosystem (Lherminier et al., 1994; Siddique et al., 1998; Kamińska et al., 2003; Christensen et al., 2004; Wei et al., 2004). The symptomatology of the disease is not solely determined by phytoplasma titre, it varies with environmental factors and plant age at the time of infection (Marcone, 2014), suggesting complex interplays with the host physiology. Phytoplasmas lack many genes of the core metabolic processes, leading to auxotrophy for many nutrients that must be supplied by the highly specialized phloem environment (van Bel and Musetti, 2019).

Among the consequences of the infection that are observed in infected plants, the most frequent are disruption of photoassimilates distribution (Musetti et al., 2013; Pagliari et al., 2016), increased or decreased sugars and starch in source or sink leaves, depending on pathosystems (Lepka et al., 1999; Maust et al., 2003; Kim et al., 2009; Tan et al., 2015; Xue et al., 2018), altered accumulation of amino acids, organic acids and secondary metabolites (Musetti et al., 2000; Choi et al., 2004; Srivastava et al., 2014; Xue et al., 2018) and impairment of photosynthetic processes (Bertamini and Nedunchezhian, 2001; Maust et al., 2003; Tan et al., 2015; Xue et al., 2018). Some of these effects have been related to callose deposition at sieve plates and aggregations of SE protein filaments, leading potentially to SEs occlusions (Hren et al., 2009; Musetti et al., 2010; Musetti et al., 2013; Santi et al., 2013).

Studies on symptomatic, well-established phytoplasma infections also reported disorganization of the vascular tissues (Oshima et al., 2001; Wei et al., 2004) and transcriptional reprogramming of genes involved in sugar transport and metabolism (Hren et al., 2009; Kim et al., 2009; Mou et al., 2013; Prezelj et al., 2016; Wang et al., 2018; Xue et al., 2018). Such effects could be triggered by phytoplasma for nutrition, plant defense response or physiological adjustments of impaired phloem activity (Christensen et al., 2005). Finally, phytoplasmas secrete effectors (Sugio et al., 2011b), that spread laterally from the SE(Bai et al., 2009; Hoshi et al., 2009; Sugio et al., 2011a). The early steps of the infection, however, are poorly known, but it would be helpful to better understand how these bacteria affect their host’s metabolism and phloem function.

The Stolbur phytoplasma - tomato pathosystem is a common model for studying plant- phytoplasma interactions. ‘*Candidatus* Phytoplasma solani’, the phytoplasma responsible for stolbur disease belongs to the 16SrXII group (Quaglino et al., 2013). Several strains infect tomato, the PO-strain causing severe symptoms, including leaf stunting, abnormal floral buds and flowers, associated with an activation of SA-mediated defense responses (Ahmad et al., 2014). Phloem hyperplasia and callose deposits are present in symptomatic leaves (De Marco et al., 2016) and phytoplasma accumulate massively into infected SEs (Buxa et al., 2015; Musetti et al., 2016). As for other plant-phytoplasma interactions, the infection affects sugar homeostasis, with alterations of sucrose synthase and invertase activities in both mature and young leaves of Stolbur-infected plants (Machenaud et al., 2007). So far phloem activity in response to the infection has not been investigated.

Sugar metabolism and phloem transport are well-documented for tomato. Sucrose loading into the phloem involves transporter-mediated sucrose transfer from the apoplasm into the SEs, so-called apoplasmic loading. Sucrose is loaded in the minor veins by Sucrose Transporter 1 (SUT1), a high-affinity sucrose proton symporter localized to plasma membrane of the SEs (Kühn et al., 2009). Unloading in sink regions requires a low-affinity sucrose transporter SUT2 (Barker et al., 2000). In midribs and on the entire length of the axial pathway from source to sink, i.e along the transport phloem, both SUT1 and SUT2 could regulate sugar release and retrieval, with SUT1 acting on sugar retrieval and SUT2 on leakage in the apoplasm (Kühn and Grof, 2010), and potentially contributing to the release/retrieval equilibrium depending on the stages of development (Minchin and Thorpe, 1987). Phloem loading fluctuates depending on the environment, with SUT1 likely involved in these regulations (Xu et al., 2018). Other classes of sugar transporters, such as sugar facilitators from the Sugar Will Eventually Be Exported Transporters family (SWEET), act on intercellular and intracellular sugar translocation (Eom et al., 2015). In tomato, the *SWEET* family constitutes a large family, with 29 members, with the genes coding for SWEET11a and SWEET12a being highly expressed in the leaves (Feng et al., 2015). Regarding sugar homeostasis, several key enzymes have been characterized, such as fructokinases (FRK), which regulate the pools of fructose and sucrose, and participate in the physiology and development of the vascular tissues (Granot et al., 2014; Stein and Granot, 2018). The regulation of sugar metabolism is expected to be highly coordinated with sugar transport. A tight link between sugar transport and metabolism implies that alterations of sugar transport in the vascular tissues can trigger modifications in phloem sap composition, which has so far not been explored.

In this study we analyzed the early responses of tomato plants infected by ‘*Candidatus* Phytoplasma solani’ to get clues on early plant response to the stolbur infection regarding phloem transport and sugar homeostasis. To determine whether the infection alters sugar long-distance phloem transport, regulated by the sucrose transporters SUT1 and SUT2, we also included in our study the two antisense lines silenced for *SUT1* and *SUT2* (Hackel et al., 2006). Altogether our results show that early plant responses to phytoplasma infection are associated with a *SUT1*-dependent perturbation of the translocation of photoassimilates in the phloem.

## Results

### Symptoms and presence of bacteria of the WT and of *SUT1* and *SUT2* antisense lines to the infection

The responses in tomato infected by ‘*Candidatus* Phytoplasma solani’ were investigated at the beginning of the symptomatic stage of infection, both in symptomatic and asymptomatic leaves, the first symptoms appearing in the younger, upper L1 leaves at the plant apex. The only visible symptoms in infected WT and *SUT2*-AS plants were beginning of yellowing, slight crook- shape and reduced growth of the L1 leaf (Fig. 2A,B,D, Table S3) and were milder than those that occur at later stages of infection (Ahmad et al., 2014). In contrast, infected *SUT1*-AS plants showed mild to no symptoms (Fig. 2C, Table S3), even though phytoplasmas were present in the L1 leaves of infected plants in all three genotypes with higher values in *SUT2*-AS (Fig. 2E,F). The L4 leaves showed no symptoms, despite the presence of phytoplasmas there (Fig. 2F). The L4 leaves of *SUT1*-AS plants had low bacterial rRNA, confirming a difference in the susceptibility of this genotype. This difference persisted during the following two weeks on *SUT1*-AS plants with weaker symptoms (Table S3). In the sixth leaf (L6), which had only just emerged at the time of grafting, only traces of stolbur rRNA were found in all three genotypes.

**Figure 1.**
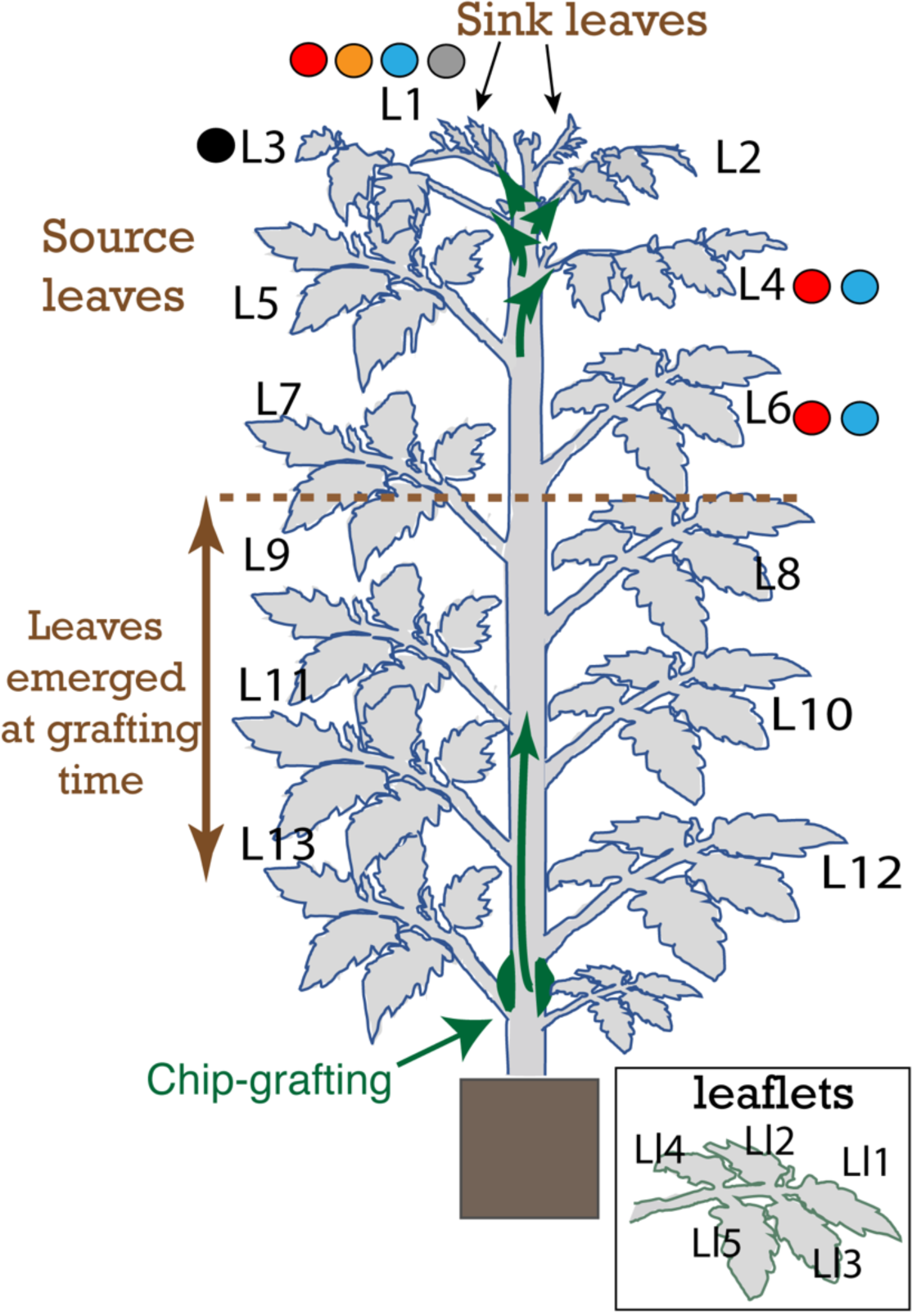
Experimental design. Schematic representation of tomato plants when material from leaves L1, L3, L4, and L6 was collected for RNA, DNA, sugar and starch analysis, sap collection by exudation and imaging. Source and sink status of the leaves follows leaf development and expansion. In this study, leaves that are more than 60% fully expanded were considered as sources, based on the study of Turgeon (1989) who established that leaves begin to export when they are 30-60% fully expanded (Turgeon, 1989). L1 leaf just emerged and began to unfold at sampling stage and was considered as a sink, with leaves L3 and older as sources, and L2 indeterminate. The arrow in green indicates the direction of migration of phytoplasmas from the grafted area to the apical leaves. Based on the age of leaves in which phytoplasma were detected at 18 days after grafting (L1to L4) and the number of leaves that had emerged after grafting (6 new leaves), it is likely that it took at least one week for graft-union to be successful and for the phytoplasma to enter the translocation stream. Samplings and observations are indicated as solid circles: plant and phytoplasma RNA sampling (red circle); phytoplasma DNA sampling (orange circle); sugars and starch sampling (blue circle); imaging by transmission electron microscopy or light microscopy (grey circle); exudate, phloem-enriched exudate sampling for metabolomics analysis (black circle). Inset showing leaflets numbering within a leaf (Ll1 to Ll5).

**Figure 2.**
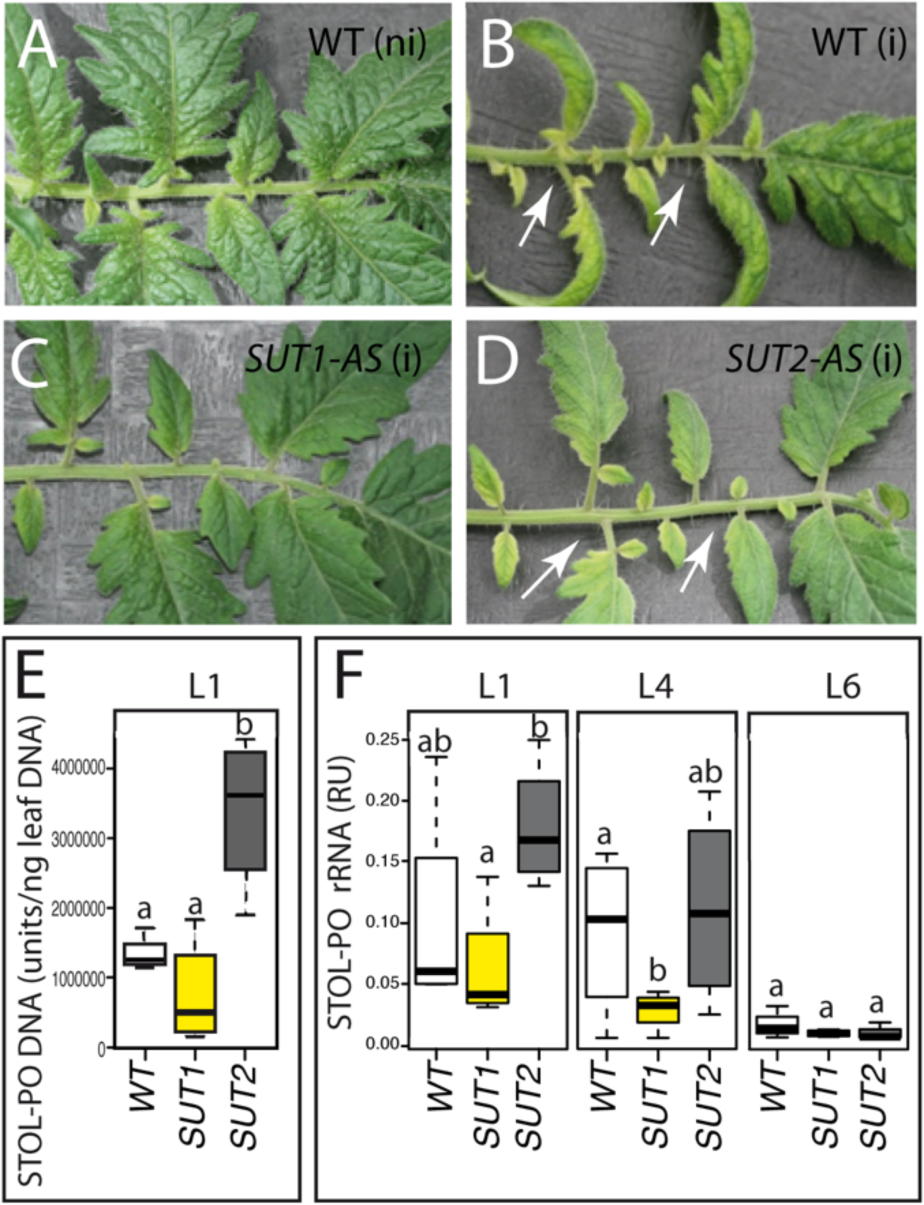
Symptoms and stolbur phytoplasma proliferation in infected plants. A to D, Details of L1 leaves from grafted non-infected WT (A), infected WT (B), infected *SUT1*-AS (C), infected *SUT2*-AS. ni: non-infected, i: infected (D). White arrows indicate leaf typical yellowing and growth reduction. E-F, Boxplot showing phytoplasma DNA in L1 leaves (E) and rRNA amounts in L1, L4 and L6 leaves; RU: relative units of content (F). The box and whisker plots in E and F show the distribution of the biological replicates. Inside black lines represent medians, top and bottom ends of the boxes represent the first and the third quartiles, respectively, and whisker extremities (open circles) represent maximum and minimum data points. White boxes: WT; yellow: *SUT1*-AS; dark grey: *SUT2-*AS; *n*=4. Different letters denote statistically different values.

### Ultrastructure of the phloem in infected leaves

We investigated the anatomy of the vascular tissues in L1 leaves, since hypertrophy of the vascular parenchyma cells characterized well-established stolbur infection (De Marco et al., 2016). In non-infected plants, the histology of the midribs was similar regardless of the genotype (Fig. S2). In infected plants, no changes were observed either, regardless of the genotype (Fig. S2A-F): neither phloem nor xylem proliferation was detected. Looking at the ultrastructure of phloem cells, we imaged at medium and high magnification 58 SE from healthy plants and 200 SE from infected plants (Fig. 3A-I). Bacteria were visible in the SEs of infected plants (Fig. 3D-I). No differences were noticed in the midrib histology of L1 leaves, when comparing WT and AS lines in either control or infected plants. Callose deposits in the SEs did not differ between healthy and infected plants, irrespective of the genotype. At a cell level, the average cross-sectional area of the SEs was less in *SUT1-* and *SUT2*-AS plants than in WT, with infection having a small additional effect. The SE area reduced to 50% or 40% of WT for *SUT1-AS*, and to 65 and 55% of WT for *SUT2*-*AS*, in healthy and infected plants, respectively (Fig. 4).

**Figure 3.**
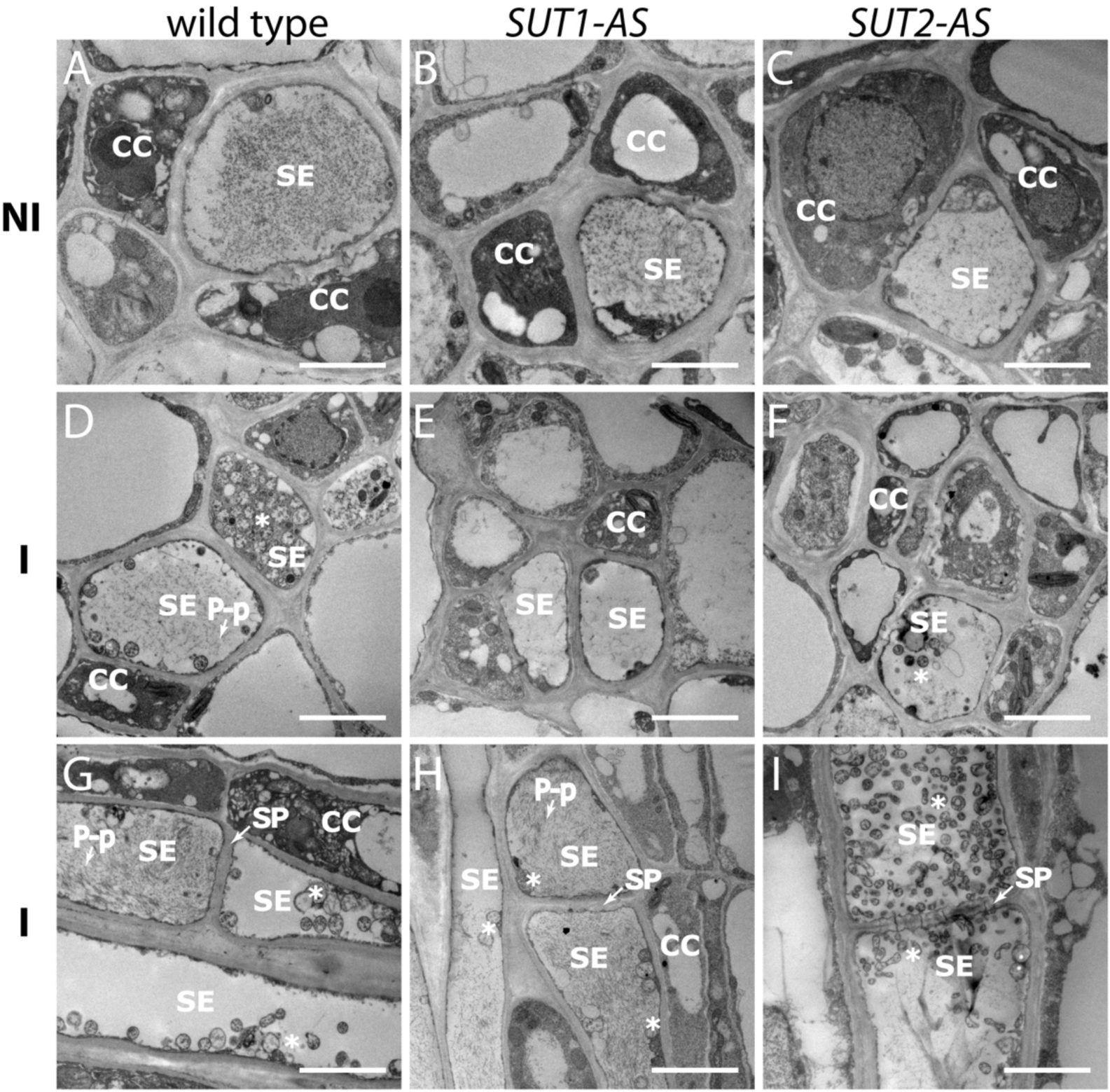
Ultrastructure of the phloem in response to the infection in the L1 leaf. Transmission electron (TEM) images of the phloem in non-infected (NI) (A to C) and infected (I) (D to I) plants. Images are representative of the sieve elements observed, with *n*=11 to 29 for healthy plants and *n*=63 to 73 for infected ones with 58 SE in total observed for healthy and 200 SE in total for infected plants. A,D,G: WT, B,E,H: *SUT1*-AS (AS), C, F, I: *SUT2*-AS plants*. A to F:* transversal sections and G to I: longitudinal sections. Phytoplasma (*) were detected in infected samples. CC, companion cell; SE, sieve element; SP: sieve plate, P-p: P-proteins. Bars, 2.5 µm.

**Figure 4.**
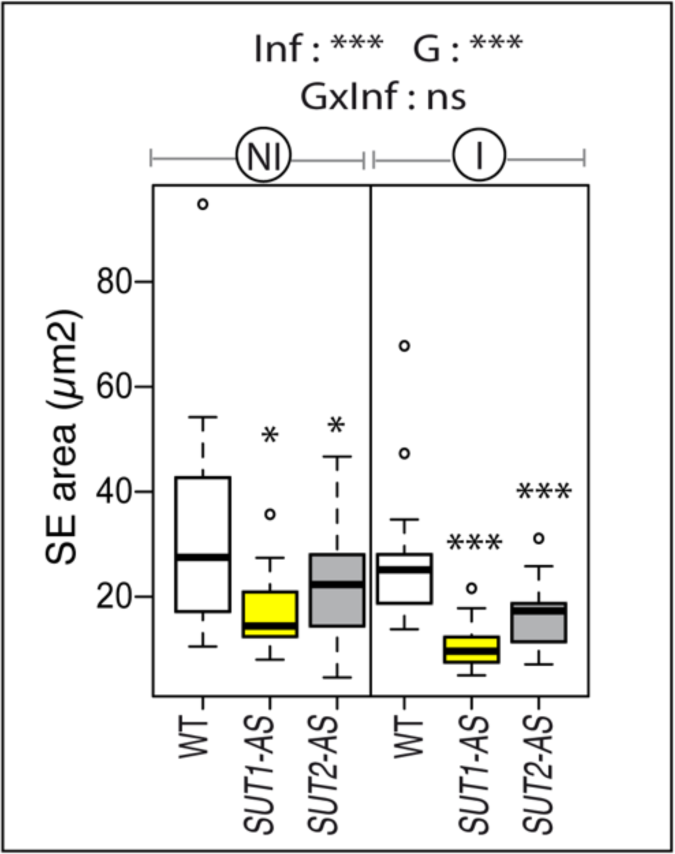
Cross section areas of SEs in WT, *SUT1*-AS and *SUT2*-AS lines. SE cross-sectional areas in the phloem of not-infected and infected plants, determined from TEM images (*n*=8 to 22). Above the boxplot, significance of the effects due to the infection (Inf), genotype (G), and their interaction (G x Inf), determined using a two-way ANOVA (*, *P* < 0.05; **, *P* < 0.01; ***, *P* < 0.001; ns, not significant). Stars above the boxes indicate significant differences by a *t*-test in *SUT1*- or *SUT2*-AS plants compared to WT plants in the same conditions (*, *P* < 0.05; **, *P* < 0.01; ***, *P* < 0.001; ns, not significant).

Phytoplasma, recognizable by their round-shaped bodies enclosing DNA strands and granular ribosomes (Fig. S3A), were abundant in the SEs of WT (Fig. 3D,G) and *SUT2*-AS plants (Fig. 3F,I), observed in most SEs imaged on of the longitudinal sections (Fig. S3B), but more difficult to identify in the SEs of *SUT1*-AS plants (Fig. 3E,H, Fig. S3B), consistent with the low rRNA (Fig. 2f). They were located either in the lumen or at a parietal location (Fig. S3C-G), although no reorganization of the plasma membrane and sieve element reticulum was observed, by contrast to later stages (Buxa et al., 2015).

We observed contacts between the plasma membranes and parietal phytoplasmas through fibrillar structures (Fig. S3C,D,F), with in some cases embedding of phytoplasmas by the sieve element reticulum (Fig. S3E). Peroxisomes, recognizable by their typical crystals, were observed in phloem parenchyma cells and at the periphery of the vascular bundles (Fig. 5A-F), with a higher frequency in infected plants, with 1 peroxisome per ROI in infected plants compared to 0.2 in non-infected plants (*p*-*value*=0.002, Fig. S4).

**Figure 5.**
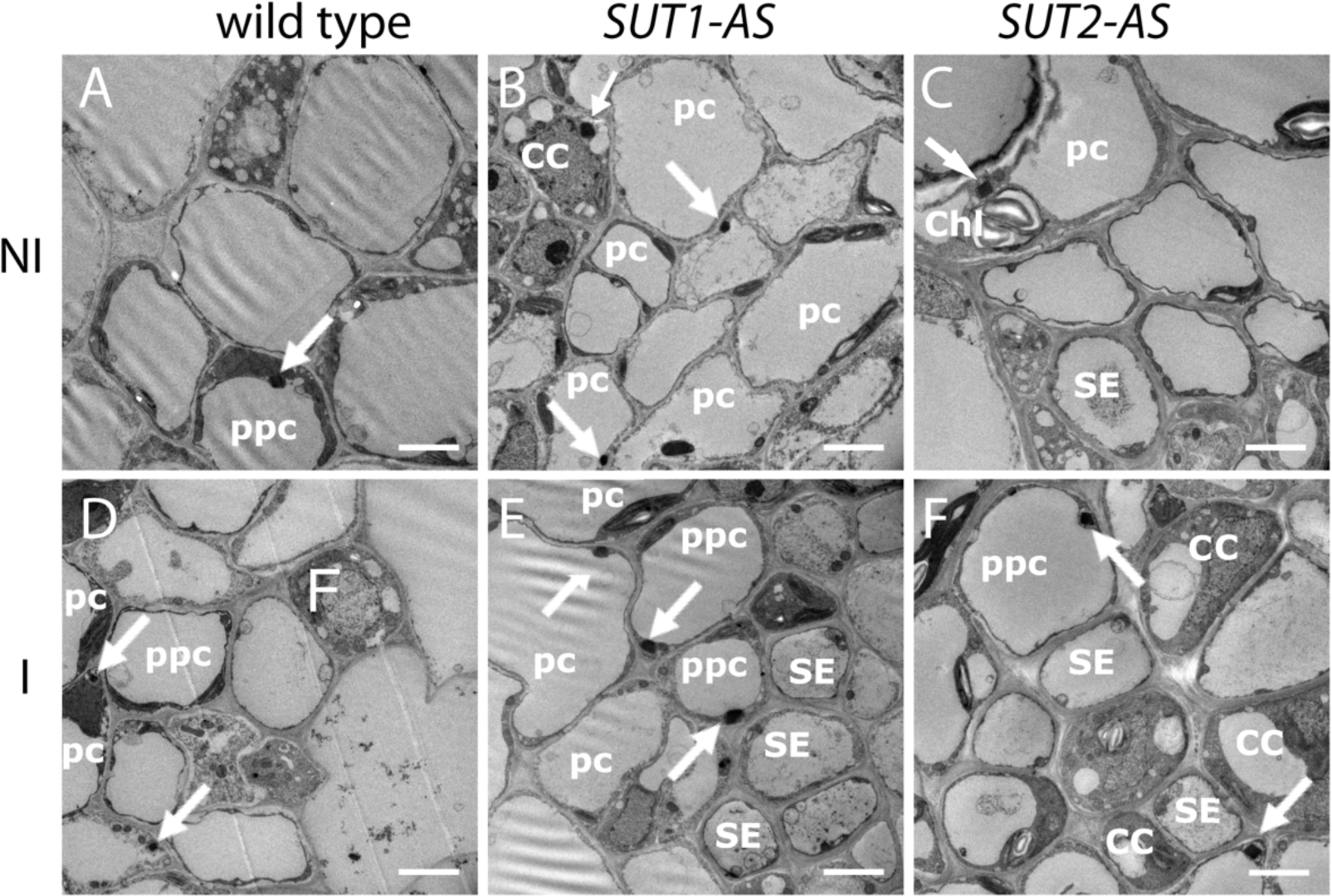
Peroxisomes in the phloem of infected and non-infected tomato plants. A to F: TEM images of the main vein phloem cells in leaf L1 in not-infected (NI) (a-c) and infected (I) (d-f) plants. (A, D) Wild-type (WT), (B, E) *SUT1*-AS antisense line (AS), (C, F) *SUT2*-AS transversal sections showing the location of peroxisomes (white arrows) in the phloem cells. Peroxisomes, easily recognizable by their large crystals, were found in parenchyma cells (pc), inside or close to the phloem bundle, in the three lines, and rarely observed in other phloem cell types (i.e.: in companion cell, in B). More peroxisomes were observed in the infected samples compared to the not-infected ones. In C, typical spatial association between a peroxisome, a mitochondrion (circle) and a chloroplast (Chl). CC, companion cell; SE, sieve element; pc, parenchyma cell; ppc, phloem parenchyma cell. Bars, 2.5 µm.

### Leaf sugar and starch content in response to the infection

Because phytoplasma infection can lead to phloem occlusions and impair photoassimilate translocation in host plants, we analyzed sugar and starch content in the lamina of L1, L4 and L6 leaves of non-infected and infected plants (Fig. S5). *SUT1*-AS plants showed higher glucose and fructose contents in the L6 leaves compared to WT plants, which is consistent with previous report on tomato plants (Hackel et al., 2006). Surprisingly, there was little effect of infection, except for a higher starch content in L1 leaves of all three genotypes, and subsequent lower sucrose-to-starch ratio. No effect was observed on the hexoses-to-sucrose ratios in L1 and L4 leaves, confirming that there was little variation in the steady state level of soluble sugars in WT and AS lines regardless of infection.

### Infection impairs phloem exudation of sugars and organic acids

Even if there was no effect of phytoplasma on leaf sugar homeostasis in mature L4 and L6 leaves, we measured in non-infected and infected plants the rate of phloem sugar exudation. It was measured on the L3 leaf, which is similar to L4 leaf for leaf expansion, both being source leaves (Fig. 6). We observed a major genotypic effect on sugar and sucrose exudation rates in non-infected plants (Fig. 6A,B), indicating that the disruption of *SUT1* and *SUT2* has impaired phloem sugar release (ANOVA *p*-value≤0.01), with an exudation rate for *SUT1*-AS plants reducing to 27 % of that for WT plants and to 56% for *SUT2-AS* plants. The L3 leaves of infected WT plants had a lower exudation rate of sugars and sucrose (38% of non-infected plants). The infection of the AS plants caused no further significant reduction than the effect of transporter disruption. Remarkably, in infected plants there was a positive correlation between the exudation rate measured in all three genotypes and the average bacterial rRNA accumulation in L1 and L4 leaves (*R*= 0.746, *p-value* = 0.0053).

**Figure 6.**
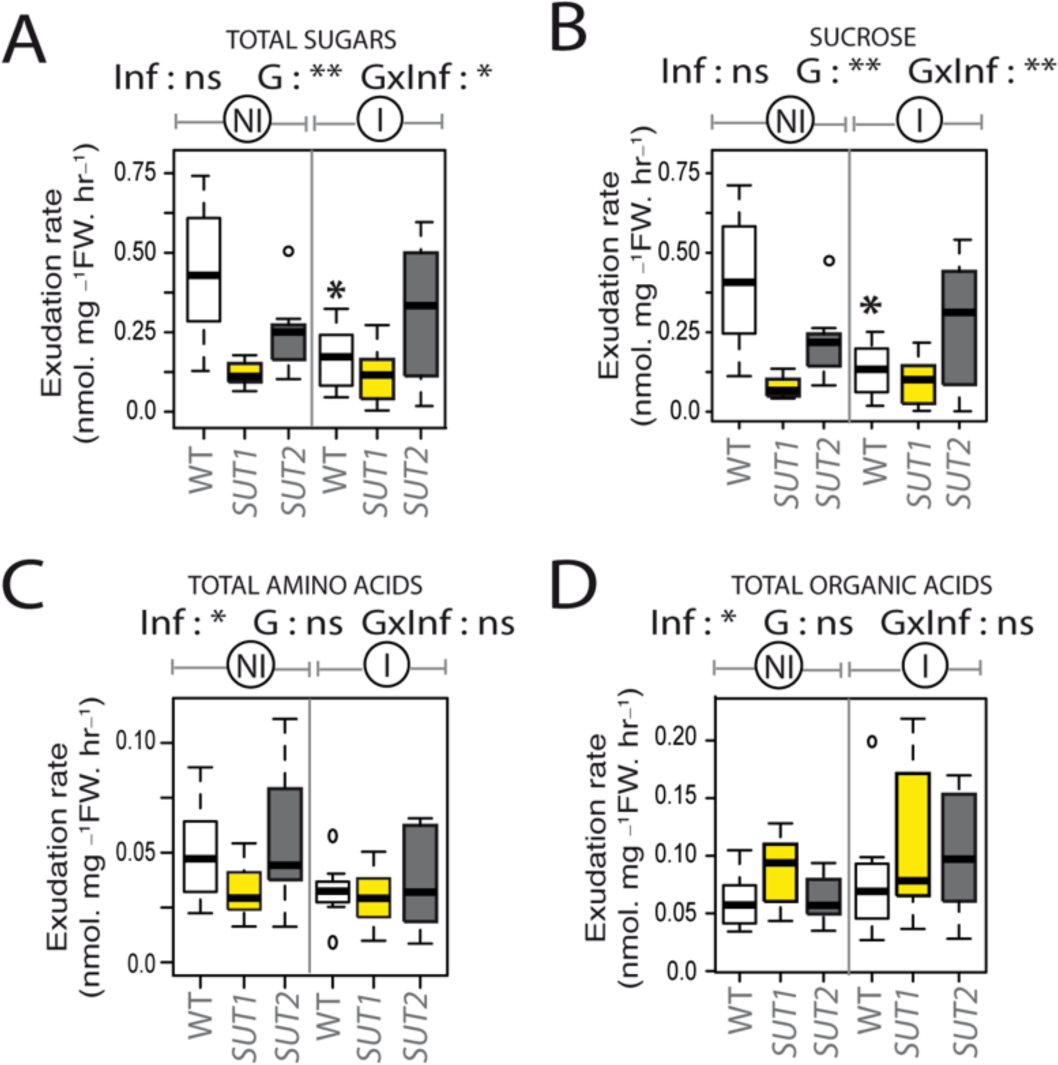
Rate of phloem exudation of metabolites from L3 leaves in response to the infection. Exudation rate is expressed in nmol mg^-1^ fresh weight (FW) per four hours of exudation. Boxplots show rates of total sugars (A), sucrose (B), total amino acids (C), total organic acids (D), in non-infected (NI) and infected plants (I). The probabilities obtained by a two-way ANOVA indicating the significance of the differences due to the Infection, to the Genotype and to their interaction Genotype by Infection, are shown on each boxplot header with “Inf” for infection effect, “G” for genotype effect, and “G x Inf” for the effect of the interaction between the genotype and the infection, with *: P < 0.05; **: P < 0.01; ***: P < 0.001; ns, not significant. The box and whisker plots show the distribution of the biological replicates. Inside black lines represent medians, top and bottom ends of the boxes represent the first and the third quartiles, respectively, and whisker extremities (open circles) represent maximum and minimum data points. White boxes: Wild-type; yellow: *SUT1*-AS; dark grey: *SUT2*-AS; *n*=7-8. Above each boxplot, significance of the effects due to the infection (Inf), genotype (G), and their interaction (G x Inf), determined using a two-way ANOVA (*, *P* < 0.05; **, *P* < 0.01; ***, *P* < 0.001; ns, not significant). Stars above the boxes indicate significant differences by a *t*-test in infected plants compared to the non-infected plants of the same genotype.

Amino acids and organic acids were also measured in the phloem-enriched exudates of non-infected and infected plants (Table **S4**). In exudates, the most abundant amino acids were glutamine, serine, asparagine, alanine and the non-proteinogenic GABA. The most abundant organic acids were malate, glycolate and glyoxylate (Table S4). No genotypic effect was observed for the exudation rates of amino acids and organic acids (Fig. 6C,D), but the ANOVA showed an reduction of the exudation rate of amino acids in infected plants (Fig. 6C). An opposite effect was observed on the exudation rates of organic acid, with higher values in infected plants compared to non-infected ones (Fig. 6D).

### Metabolite content of phloem-enriched exudates

The lower sucrose exudation rate observed for infected WT plants could result from sugar consumption by bacteria or from cleavage of sucrose in the SEs to provide precursors for the synthesis of callose or cell wall precursors. To identify, within the metabolite profiles of the phloem-enriched exudates, specific differences in their proportion, independent of the exudation rate, the profiles were adjusted using a method of normalization (Fig. S1) that has been developed for the analysis of phloem-enriched exudates (Yesbergenova-Cuny et al., 2016). The normalized values, termed “content”, showed high positive correlations between infected and non-infected metabolite profiles (*R^2^* > 0.97, Fig. 7A), revealing a strong homeostasis in phloem-enriched exudate composition. In non-infected plants there was no modification of the sucrose content in the L3 exudates of the three genotypes (Fig. 7B), yet glycolate content was higher (Fig. 7B). The content of branched amino acids (valine, leucine and isoleucine), aspartate, glutamine, proline, serine and glycine, also varied with genotype (Fig. 7B). In response to phytoplasma infection, the content of most metabolites was not altered (Table S5, Fig. S6). Malate content, an abundant organic acid, showed no variation. Nevertheless, irrespective of genotype, infection slightly reduced content of sucrose (10% less compared to non-infected) (Fig. 8A). Infection increased content of glycolate and aspartate in L3 exudate of WT plants (approximately 80 and 50% more, respectively) (Table S5), and of glyoxylate in both AS genotypes (approximately two-fold increase) (Fig. 8B,C, Table S5). No correlation was observed between the content of the exudate metabolites and the average bacterial rRNA accumulation quantified in L1 and L4 leaves.

**Figure 7.**
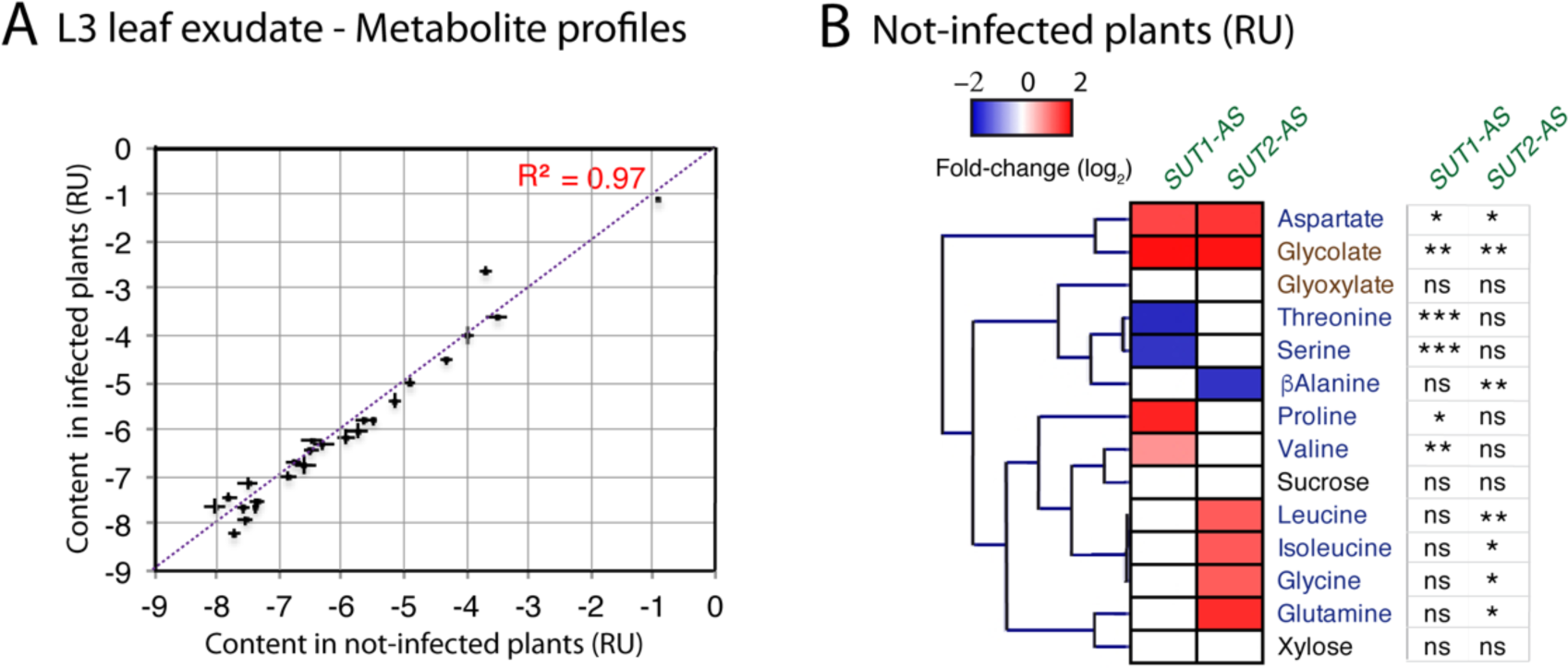
Comparison of metabolite profiles of the phloem-enriched exudate from the L3 leaf of not- infected and infected plants. The metabolite relative content was determined on the exudate of the third leaf for wild-type (WT), *SUT1-* and *SUT2*-AS lines. A: Pairwise comparisons and R^2^ correlation coefficient between metabolite profiles in infected and not-infected plants in the different genotypes and conditions. The plot show for each metabolite its content in the exudates of not-infected plants (X-axis) and infected plants (Y-axis). A linear regression has been drawn on the plot, which showed that most metabolites remained stable in both conditions, illustrating homeostasis of the most abundant primary metabolites in the phloem- enriched exudates. B: Heat map showing significant fold-changes in metabolite content in the phloem- enriched exudates from the L3 leaves of not-infected *SUT1*-AS and *SUT2*-AS plants compared to not- infected WT plants (*n*=7-8). Values are shown in a blue-to-red log2 scale with blue for negative values, red for positive values and white for no difference. On the right panel: significance of the effects due to genotype (G), determined using a one-way ANOVA (*, *P* < 0.05; **, *P* < 0.01; ***, *P* < 0.001; ns, not significant).

**Figure 8.**
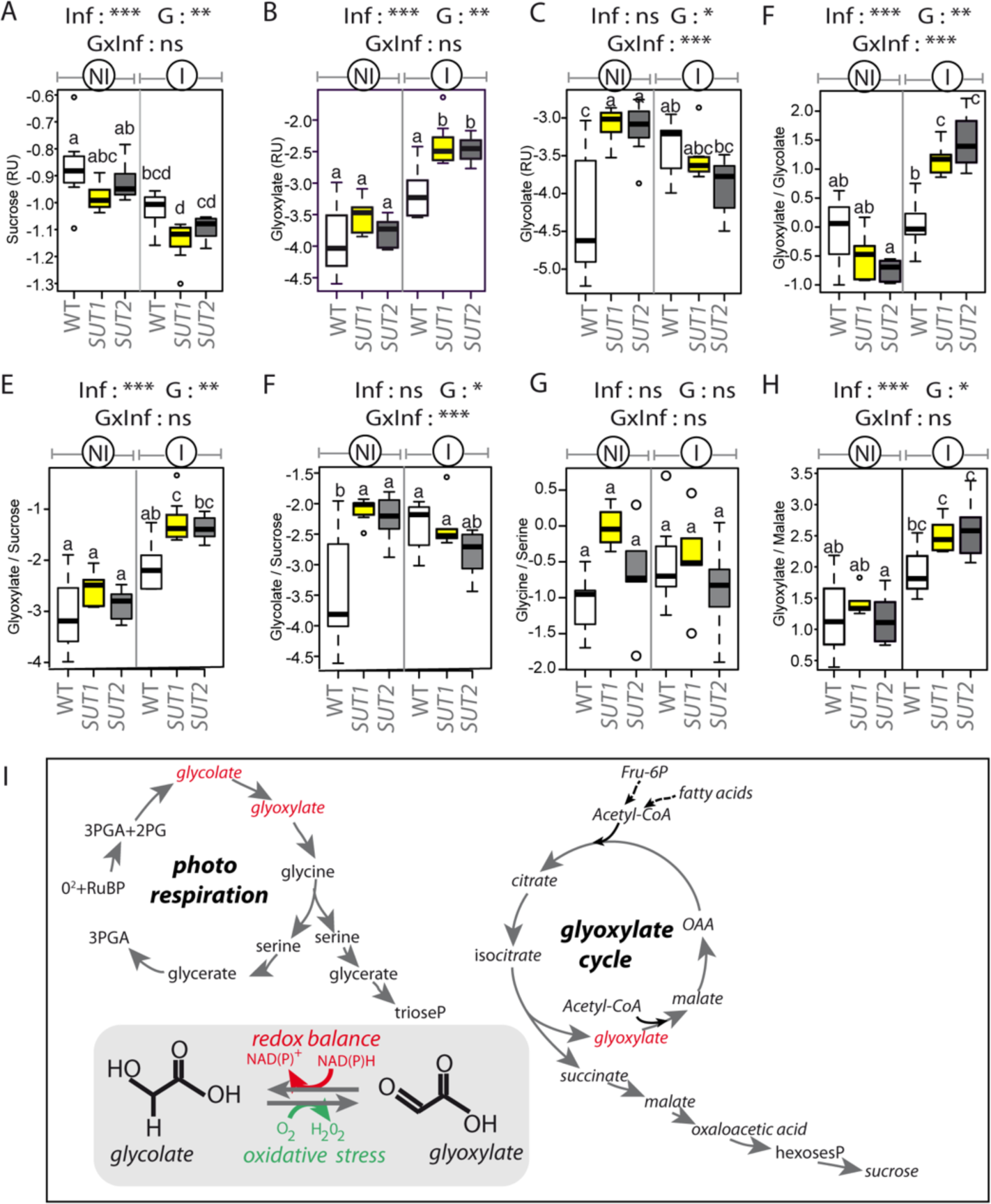
Main variations in the metabolite content of phloem-enriched exudates in response to the infection. A to H: Boxplot with the content in the exudate from the L3 leaf of (A) Sucrose, (B) glyoxylate (C) glycolate; and the content ratios for (D) glyoxylate-to-glycolate, (E) glyoxylate-to-sucrose, (F) glycolate-to-sucrose, (G) glycine-to-serine, (H) glyoxylate-to-malate ratios. White boxes: Wild-type, yellow: *SUT1*-AS; dark grey: *SUT2-*AS, *n*=6-8. Non-infected plants: NI, Infected plants: I. RU: relative units for content, plotted on a log2 scale. The box and whisker plots show the distribution of the biological replicates. Above each boxplot, significance of the effects due to the infection (Inf), genotype (G), and their interaction (G x Inf), determined using a two-way ANOVA (*, *P* < 0.05; **, *P* < 0.01; ***, *P* < 0.001; ns, not significant). (i) Photorespiratory pathway and its interaction with the glyoxylate cycle. Glyoxylate can be produced either via the photorespiratory pathway or via the glyoxylate cycle, the latter being a bypass of the TCA cycle in conditions of carbon limitation, glycolate and glyoxylate being inter-converted depending on the redox balance (Dellero et al., 2016). 2-PG, 2 phosphoglycolate; 3-PGA, 3 phosphoglycerate; RuBP, ribulose 1,5-bisphosphate. Glyoxylate and glycolate are reversibly converted by glyoxylate reductases (GLYR) and glycolate oxidases (GOX), reactions that are controlled by the redox status and contribute to the production of reactive oxygen species (ROS) and the conversion of NADPH into NAD(P)^+^.

### Variations in glycolate and glyoxylate in phloem-enriched exudates

According to the ANOVA, infection explained most of the variation in sucrose and glyoxylate in the exudate (*Pinfection*<0.001) (Fig. 8A,B), while glycolate was strongly affected by the interaction “infection per genotype” (*Pinfection x genotype*<0.01) (Fig. 8C). The variation of the glyoxylate-to-glycolate ratio was explained by the effects of the infection, of the genotype and the interaction “infection per genotype” (Fig. y). There was a strong effect of infection on the glycolate-to-sucrose ratio (Fig. 8E) and glyoxylate-to-sucrose ratio (Fig. 8F). Slight variations in the glycine-to-serine ratio, a marker of photorespiration (Foyer et al., 2003), were not explained by the genotype or the infection (Fig. 8G). For the glyoxylate-to-malate ratio, a strong effect of the infection was observed (*Pinfection*<0.001) (Fig. 8H).

### Transcriptional reprogramming of selected genes

The above-mentioned hypotheses were further investigated by the analysis in L1, L4 and L6 leaf midribs of the expression of genes encoding either stress markers or involved either in glyoxylate cycle, photorespiration or sugar transport and metabolism (Tables S1 and S2). The genes selected from available information in tomato databases (Toufighi et al., 2005; Goodstein et al., 2012) were coding for glycolate oxidases (*GLO00/GOX2*, *GLO40/GOX1*, *GLO50/GOX3*), glyoxylate reductases (*GLYR1* and *GLYR2*), isocitrate lyase (*ICL*) and malate synthase (*MLS*). Genes coding for callose synthases (*CAS2* and *CAS7*) and PR proteins (*PR1a* and *PR2a*) were included as stress markers. *CAS7* was reported to be the only callose synthase gene, out of eight retrieved from the NCBI database, to show an increase expression in symptomatic tomato leaves in comparison to non-infected ones (De Marco et al., 2016). For sugar transport and metabolism, we included genes coding for fructokinases (*FRK1*, *FRK2*, *FRK3*), sucrose synthases (*SUS1* and *SUS3*), *SWEET* sugar facilitators (*SWEET2a, SWEET5b, SWEET10c*, *SWEET11a*, *SWEET12a*). A gene coding for a phloem marker (*Phloem protein 2*, *PP2*) was added as well. The expression of most genes varied depending on the leaf (Table S6) so the responses were analyzed per leaf level (Fig. 9). A strong correlation (*R*>0.7, *p*value < 0.001) was found between the expression of *CAS7*, *PP2* and *FRK3* (Table S7).

**Figure 9.**
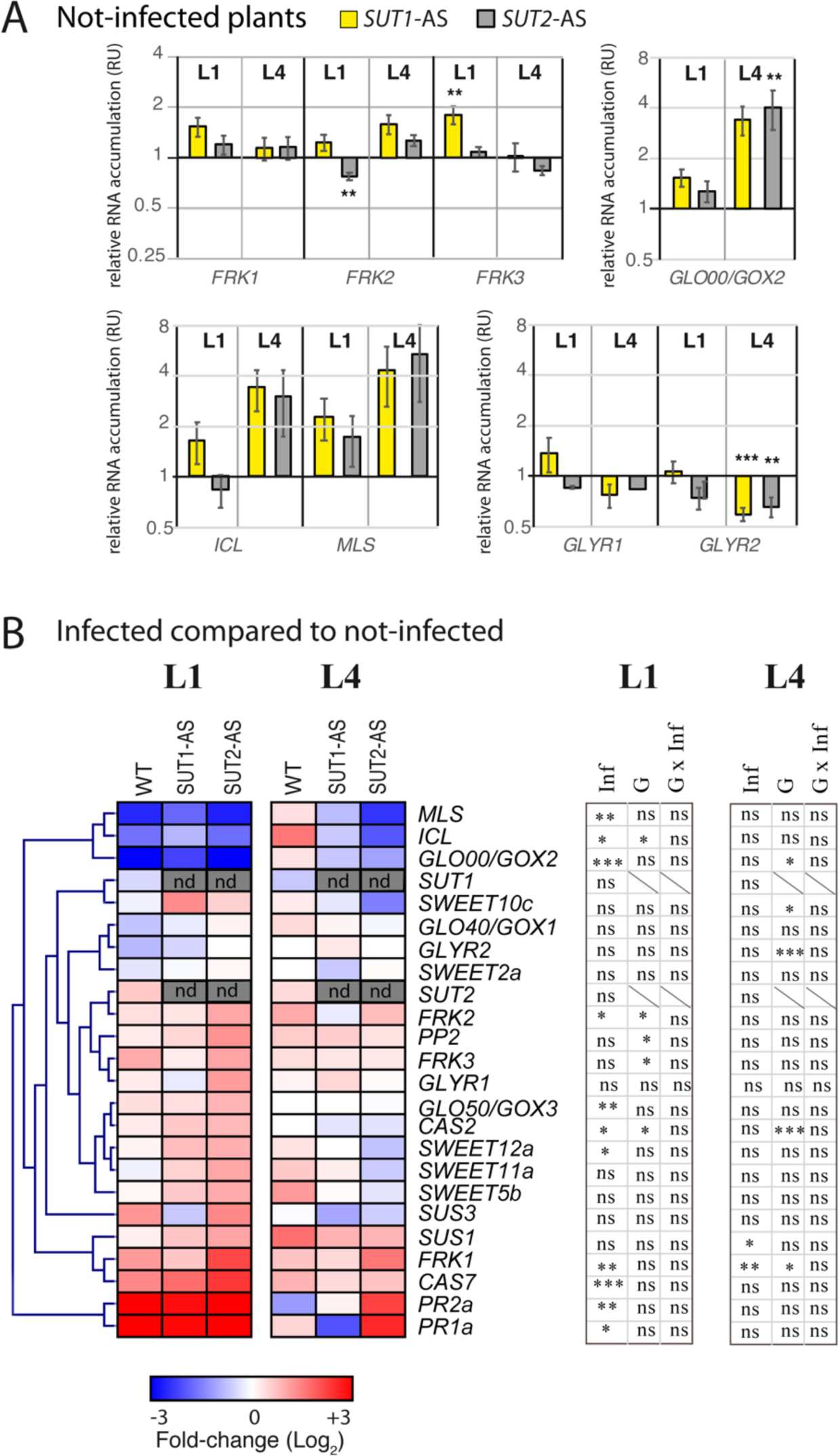
Transcript profiling of candidate genes in not-infected and infected plants. A: Transcript profiles of genes in not-infected *AS* plants compared to non-infected wild-type (WT) plants, with mRNA in L1 and L4 leaves. The accumulation of transcripts was normalized by the mean accumulation in WT plants. Histograms show for each gene mean response +/- *SE* (*n*=4). Yellow bars: *SUT1*-AS plants; Dark grey bars: *SUT2*-AS plants. Y-axis: Relative transcript accumulation, reported to the mean value of WT plants set to 1, Y-axis is drawn with a log2 scale. RU: relative units for content, *t*-test: *: P < 0.05; **: P < 0.01; ***: P < 0.001; ns, not significant. B: Inserted heat maps showing fold-changes between infected and non-infected plants in each genotype in L1 and L4 leaves (*n*=4). Fold changes were determined after normalization by the reference genes; values are shown on a log2 scale, with blue values for metabolites showing a smaller content due to infection, and red values for higher content. In grey, not determined (nd). On the right side, results of the two-way ANOVA on expression data in the L1 and L4 leaves, showing the significance of the effects due to the infection (Inf), genotype (G) and their interaction (G x Inf) (*, *P* < 0.05; **, *P* < 0.01; ***, *P* < 0.001; ns, not significant).

In non-infected plants, differences were observed in the transcript levels for *FRK2* and *FRK3* in L1 leaves and for *GLO00/GOX2* and *GLYR2* in the L4 leaves in *SUT1*- and *SUT2*-AS plants, compared to WT plants (Fig. 9A). The data indicated that the downregulation of *SUT1* and *SUT2* affected sugar metabolism and photorespiration. In infected plants all L1 leaves showed symptoms and, correspondingly, infection raised *PR1a* and *PR2a* transcript level in all genotypes (Fig. 9B). Higher transcript levels were also observed for *CAS7*. We observed no significant response in L1 leaves on the expression of *SUT1, SUT2*, *SWEET2a*, *SWEET11a*, *SWEET5b* and *SWEET11c.* By contrast, transcript levels for *GLO50/GOX3*, *SWEET12a*, *FRK1* and *FRK2* were higher, and lower for *GLO00/GOX2*, *ICL* and *MLS* (Fig. 9B). The effects of infection were lower in L4 than L1 (Fig. 9B), with higher transcript levels of *SUS1* and *FRK1*. Interestingly, no significant changes were observed in the L4 and L1 leaves for the transcription levels of *SUT1* and *SUT2* in infected WT plants compared to non-infected. No correlation was observed between the accumulation of *PR1a, PR2a, CAS2* and *CAS7* and the accumulation of phytoplasma rRNA in L1 and L4 leaves. Remarkably, we observed several correlations between the expression of *FRK1*, *FRK3*, *SUS1* and *SUS3* and the expression of *ICL*, *MLS*, *GOX1*, *GOX2*, *GOX3* and *GLYR1* (Table S7). A negative correlation was found between the expressions of *FRK1* and *ICL* and *GLYR2 (p*value*<0.001)* while a positive correlation was found between the expression of *SUS1, SUS3* and *FRK3* and the expression of markers of peroxisomal activity (*R*>0.44, *p*value*<0.001*) (Table S7).

## Discussion

### *SUT1* and *SUT2*-AS lines as a tool to study the plant response to phytoplasma infection

Phloem loading, release or retrieval, are expected to be altered in *SUT1-* and *SUT2*-AS lines, with potential consequences on phloem flow and sap composition. In non-infected plants, the sugar exudation rate from excised leaves was decreased in the two lines compared to WT, with a marked effect in *SUT1-AS* plants, consistent with the roles of SUT1 and SUT2 in sugar transport (Hackel et al., 2006). Because both lines were affected, it is likely that the equilibrium of release/retrieval of sucrose along the transport phloem changed, and not only (re)-loading, since that is due solely to SUT1, or release, potentially due to SUT2. Unexpectedly, sucrose content was not impaired in the exudate of these lines, revealing a tight homeostasis in phloem-enriched composition, and indicating that the differences in sugar exudation rates were not due to variations in the sucrose composition of the exudate. The findings are consistent with the report of tight homeostasis of sucrose concentrations and sap osmotic potentials in *Sonchus oleraceus* during phloem pathway blockage (Gould et al., 2004). Besides, very few metabolites changed, except for higher glycolate and aspartate in the exudates of the AS lines (Fig. 7). Further study is needed to determine how the impairment of sugar transport is associated with the accumulation of aspartate and glycolate in CCs and their entry to SEs.

Phloem mass flow, being convective, is the product of sap concentration, flow velocity and cross-sectional area of functional SEs. Strikingly, the cross-sectional area of SEs in the apical leaves of *SUT1-* and *SUT2-*AS plants was reduced compared to WT. Together, our data suggest that the decrease in sucrose transport in the AS lines, may be related to decrease in SE cross-section or reduced flow velocity. Since the phloem flow is also known to reduce in infected plants (Pagliari et al., 2017), we included the AS genotypes in our study, in order to comparing the various outcomes and shed more light on effects of phloem sap composition and phloem flow on responses to phytoplasma.

### Phytoplasma accumulation, symptoms and triggering of SA-defense responses

We analyzed the responses of symptomatic and asymptomatic leaves with active phytoplasma multiplication. One day after first symptoms appeared, the bacteria had multiplied in the four apical leaves. A delay for the graft to be effective and permit propagation of the phytoplasma from the grafted zone could explain why no phytoplasma were detected in the L6 lower leaf. Surprisingly, symptoms were restricted to the L1 leaf, associated with an increase in its starch accumulation (Fig. 10) but with no anatomic alteration. The lack of symptoms in L4 leaves, where phytoplasma infection was confirmed by the detection of bacterial rRNAs, may be related to differences of leaf development or phytoplasma titers when phytoplasmas spread. A more massive invasion may have occurred for L1 sink leaf because more phytoplasmas were translocated from the graft itself, or from the lower infected source leaves. In the symptomatic L1 leaf, yellowing and slight curling were associated with an upregulation of *PR1a* and *PR2a*, both markers of the SA-signaling pathway that is induced in infected tomato (Ahmad et al., 2014). The callose synthase gene, *CAS7*, was also upregulated, which is consistent with callose deposits and *CAS7* upregulation occurring in later phases of tomato stolbur infection (De Marco et al., 2016). However, symptoms appearance and SA-mediated responses were not directly related to phytoplasma RNA accumulation in the L1 and L4 leaves. Other reports have described variations in the symptomatology (Wei et al., 2004; Marcone, 2014) that may be related to plant developmental stage or nutritional status at the time of infection. Remarkably, the *SUT1-AS* line was much less susceptible to the infection, compared to the two other genotypes, with decreased phytoplasma multiplication and milder symptoms. Such a tolerance could be the consequence of metabolic changes, supported by changes in the expression of *FRK3* or *GLYR2* (Fig. 9), to such an extent that defense cascades are being triggered more efficiently in the *SUT1-AS* line.

**Figure 10.**
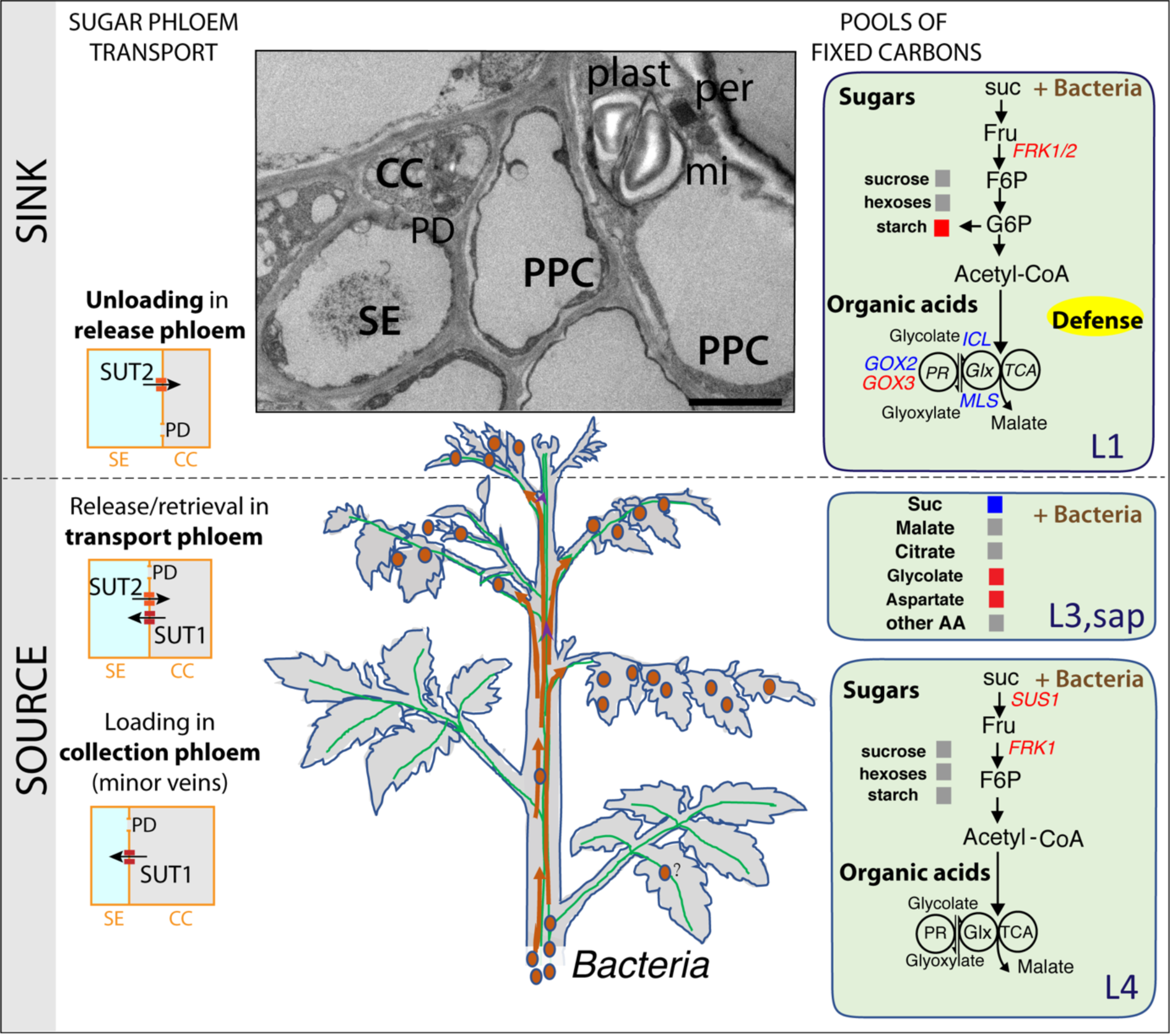
A model of our inferred responses in the young (sink) and mature (source) leaves of a tomato plant that were analyzed after infection by phytoplasmas. On the right side, primary metabolic steps regulating the levels of sugars and organic acids (main pools of fixed carbon) in L1 and L4 leaves. Upregulated genes are shown in red, downregulated genes are shown in blue. The metabolites in exudate from L3 are shown in grey (no change), blue (decrease) or red (increase). Similarly, soluble sugars and starch contents are shown in L1 and L4 leaves. The enzymes encoded by these genes act on glycolysis, photorespiration and glyoxylate cycle. Cytosolic sucrose cleavage by sucrose synthases (SUS) produces UDP-glucose and fructose, with fructose being phosphorylated by fructokinases (FRK) as a first step of glycolysis. Isocitrate lyase (ICL) and malate synthase (MLS) are the main enzymes of the glyoxylate cycle (Glx). Glyoxylate and glycolate are reversibly converted by glyoxylate reductases (GLYR) and glycolate oxidases (GOX), which also acts on the photorespiration (PR). Orange arrows: systemic transport of phytoplasma to leaves at an early stage of development. On the left side are reported the three main steps of sugar transport in the phloem with the relative contribution of the sugar transporters SUT1 and SUT2 in this process. Transmission electron microscopy inset shows phloem cell types in which take place the metabolic pathways: mitochondria for TCA cycle, peroxisomes for glyoxylate cycle and photorespiration, plastids for starch accumulation. Key: suc, sucrose; Fru, fructose; F6P, fructose-6- phosphate; Glc, glucose; G6P, glucose-6-phosphate; AA, amino acids; Glx, glyoxylate cycle; TCA, tricarboxylic acid cycle; PR, photorespiration. CC, companion cells; SE, sieve elements; PPC, phloem parenchyma cells; plast, plastid; mi, mitochondria; per, peroxisome; PD: plasmodesmata. Bar: 2.5 µm.

### Phytoplasma transport, sugar exudation rate and exudate composition

Sugar exudation rate is expected to reflect sucrose loading and export from a leaf before excision, despite any concerns arising from excision and contamination during phloem sap collection. There was a positive correlation between the rate of exudation from the L3 leaves and the phytoplasma rRNA accumulation, at least at this stage of infection. On the other hand, no significant correlations were found between metabolite contents in the exudate and bacterial rRNA accumulation, suggesting that the contents of the metabolites identified in this study are not determinants for phytoplasma multiplication. However, the first symptoms appear all at once in the three lines (Table S3), arguing that variations in the multiplication may not solely be due to differences in the kinetics of propagation along the plant. The results indicate that the number of bacteria units delivered into the translocation stream might be also reduced, which could be an alternative hypothesis to explain the low susceptibility of the *SUT1-AS* plants, and the very few bacteria in the SE of their L1 leaves, besides metabolic changes. The speeds of spread of phytoplasma are too low for the movement to be simple convection in the flow, which may indicate that the bacteria are stationary in the SEs throughout part of their lifetime, possibly anchored to the membranes. Our findings support the notion of a restriction of passive translocation due to the retainment of bacteria in source leaves, scenario that was also supported by report on the phytoplasma colonization of *Euphorbia pulcherrima* (Christensen et al., 2004).

Because phloem sugar transport has seasonal variations, our observation could help to explain the seasonal fluctuations in the colonization of fruit trees by phytoplasmas (Marcone, 2014). In addition, environmental clues, such as mineral deficiency, water deficit, low temperatures or variations in light intensity can modify phloem sugar transport (Lemoine et al., 2013; Dannoura et al., 2018; Xu et al., 2018; Epron et al., 2019), with stress conditions leading in some cases to higher carbon allocation to sink organs (Durand et al., 2016). These observations indicate that agricultural practices reducing mass flow in field conditions may be a way to limit the multiplication of phytoplasma and control disease development, with a need to balance benefit against lost production.

### What factors are necessary for the decline of exudation by the infection?

Several studies report that phytoplasma infection alters carbon allocation, related to an impairment of phloem loading, and limit bacterial infection (Christensen et al., 2005). The reduced exudation from excised source leaves supports the suggestion of Christensen et al., (2004) that reduced export may occur with infection, and help explain the lack of spread in their system to sink tissues. SE occlusion by callose deposition, aggregation of P proteins, and cell wall thickening have been proposed to partly explain the impairment carbon allocation (Lherminier et al., 2003; Musetti et al., 2013; De Marco et al., 2016), although direct evidence of their role is still pending (Pagliari et al., 2017). A reduction in phloem flow has been quantified in Arabidopsis plants infected by *Chrysanthemum yellows* phytoplasma, using carboxy-fluorescein fluorescence assay (Pagliari et al., 2017). Our data are consistent with this study, with a reduction of sugar exudation rate to one third of the level observed in non-infected plants, similar to the reduction observed in Arabidopsis. Amazingly, the overall metabolite profiles of the exudate of WT plants was similar in infected and non-infected plants, indicating a reduction in mass flow, rather than a massive reduction of sucrose concentration of the sap per se, when sap composition would have changed.

Although infection of WT plants reduced exudation from the L3 leaf dramatically, infection hardly affected exudation for SUT1 and SUT2 plants. Altogether our findings indicate that wild type plants reduce mass flow in response to the infection, while the mutants are unable to do so. If the occlusion of the sieve pores by callose depositions explain part of the reduction in mass flow, as proposed elsewhere (Lherminier et al., 2003; Musetti et al., 2013; De Marco et al., 2016), a similar response is expected to be observed in the antisense lines, which was not the case. Moreover, callose deposits did not differ in healthy and infected plants in the phloem of L1 leaves at this stage of infection, suggesting that callose deposits is likely not necessary for alteration of mass flow at this stage. Meanwhile a strong correlation was observed between expression of *CAS7* and the expression of *PP2* and *FRK3*, all three being expressed in the vascular tissues. Tomato *CAS7* is the ortholog of the Arabidopsis *CalS7*, a gene expressed in the phloem and necessary for phloem development (Barratt et al., 2011; Xie et al., 2011). The higher *CAS7* transcript accumulation may indicate a defense response, to limit the propagation of phytoplasma, but it could also reveal the onset of phloem hyperplasia that has been observed in tomato plants at more advanced stage of infections (De Marco et al., 2016), consistent with the higher transcript accumulation of *FRK3* and *PP2*. If the reduction of mass flow is explained by impaired sugar loading and release/retrieval balance, associated responses such as modifications of the phloem-enriched exudate composition are expected to be observed, what we actually observed. Our findings provide evidence that *SUT1* and *SUT2* are necessary for the decline in phloem mass flow.

### What are the consequences of the infection on the metabolism?

A slight decline in exudate sucrose content was observed in infected compared to non-infected plants, regardless of the genotype (about 10 % reduction), suggesting that this decline was independent of *SUT1* and *SUT2* functions, and that in infected source leaves, a consumption of sucrose due to phytoplasma in SEs may reduce sucrose content, a cleavage of sucrose to provide precursors for the synthesis of callose or cell wall components, or that there was a reduction of sugar loading or retrieval because of limited sucrose in the apoplasm. This latter hypothesis is consistent with the observation in source (L4) leaves of infected WT plants of higher levels of *SUS1* and *FRK1* transcripts, suggesting more sucrose is cleaved and available for glycolysis. That brings us to the question of whether the infection modifies the availability of other carbon sources in the phloem sap. Phytoplasma, having small genomes, are auxotroph for many nutrients, with malate being potentially their main source of carbon (Christensen et al., 2005; Kube et al., 2012; Saigo et al., 2014). However, infection did not affect malate content in phloem-enriched exudates. It would be worthwhile to determine whether the delivery of malate to phytoplasmas occurs via apoplasmic or a symplasmic pathways. Malate accumulates in different subcellular compartments, including the apoplasm (Lohaus et al., 1995; O’Leary et al., 2016). Close abutting of phytoplasma to the plasma membrane of the SEs at our early stage (Fig. S3), as also observed at later stages (Buxa et al., 2015; Musetti et al., 2016), suggests that some nutrients are taken from the apoplasm through connections with the SE plasma membranes.

The response to infection might be slightly different in sink leaves. In the symptomatic L1 leaves, we observed an upregulation of both *FRK1* and *FRK2* compared to non-infected L1. Both genes are expressed in the CC and in xylem cells (Stein et al., 2018). FRKs have been proposed to play a role for the supply of carbon for starch accumulation in fruit development (Schaffer and Petreikov, 1997; Stein and Granot, 2018), which may correspond to the accumulation of starch in infected L1 leaves. FRKs have also been proposed to maintain the balance between sucrose degradation and synthesis, working together with SUSs, to maintain sink strength (Davies et al., 2005; Stein and Granot, 2018). Our data showed correlations between *FRK1* and *FRK2* transcript levels with those of *SUS1* and *SUS3* (Table S7), which indicate that they could also contribute to maintain sink strengths in infected sink leaves.

Overall our findings indicate that plants react to the infection by a modification of sugar release and retrieval in the phloem, and of sugar homeostasis in the source and in the sink tissues. As a consequence, the intercellular translocation of sugars in the vascular tissues could also change, with the involvement of SWEET facilitators (Aubry et al., 2019). Sugar availability in the apoplasm would respond, as observed for other cases of plant pathogens interactions (Eom et al., 2015; Pommerrenig et al., 2020). However, we found that four of the *SWEET* genes highly expressed in the leaves (Feng et al., 2015) were not responsive to the stolbur phytoplasma, except for a slight upregulation of *SWEET12a* in L1 leaves.

What are the consequences of altered *SUT1* and *SUT2* expression on metabolism?

An unexpected result was the finding of elevated glycolate contents in phloem-enriched exudates of L3 leaves, both in the non-infected AS plants and in response to the infection in WT plants. Glycolate is an intermediary product of photorespiration and the reduced form of glyoxylate, that can be produced by the glyoxylate cycle, the two pathways being located in peroxisomes (Hu et al., 2012). Glycolate and glyoxylate are interconverted by glycolate oxidases (GOX) or glyoxylate reductases (GLYR). Peroxisomal metabolism is involved in many metabolic pathways, regulating the redox homeostasis, production of ROS and synthesis of hormones required in plant immunity (Igamberdiev and Eprintsev, 2016; Yuan et al., 2017; Kao et al., 2018). Several reports indicate that the glyoxylate cycle, producing succinate and malate, can be induced when plants are responding to pathogens and phloem-feeding insects, a response that may contribute to carbon reallocation (Cots et al., 2002; Bolton et al., 2008; Peng et al., 2016). The higher levels of transcripts of *GOX2,* and the low levels of transcripts of *GLYR2*, in the source leaves of the antisense lines further confirmed a metabolic switch related to peroxisomal activity when *SUT1* or *SUT2* expression was impaired. Interestingly, positive and negative correlations were found between the transcript levels *FRK1, FRK3* and *SUS3*, and those of *GOXs*, *GLYRs* and *ICL* (Table S7), supporting our hypothesis of an association between sugar homeostasis and peroxisome metabolism. We should now consider how these metabolic effects, which could be related to sugar transport impairment, will affect redox homeostasis. Several studies report that redox status plays a role in the regulation of SUT transporters, of sugar phloem loading (Krügel et al., 2008; Asensi-Fabado et al., 2015) and of the activities of SUSs and FRKs (Dumont and Rivoal, 2019; Farooq et al., 2019). The contribution of peroxisome metabolism to regulating phloem sugar transport needs attention.

## Conclusions

The goal of this study was to compare the responses to phytoplasma infection in symptomatic and asymptomatic leaves and to evaluate the role of phloem sugar transporters in the reduction in sugar export triggered by the infection. Absence of a correlation between phloem-enriched exudate composition in abundant primary metabolites and phytoplasma multiplication indicates that phloem sap composition is not limiting for the infection cycle. The reduction in the exudation rate that occurs from infected source leaves, and the metabolic changes in sap composition associated with *SUT1* and *SUT2* expression, indicate that *SUT1* and *SUT2* are engaged in this process. Finally, our findings do not support the hypothesis that callose deposition is the main process reducing phloem mass flow in response to phytoplasma infection.

## Materials and methods

### Plant material and infection by phytoplasma

Tomato plants (*Solanum lycopersicum* L., cv. Money Maker) were grown in glasshouse (27°/20° C day/night, in a 16h photoperiod), in soil with sand and organic matter (20:80). Seeds of the antisense (AS) lines *SUT1-15* (hereafter called *SUT1*) and *SUT2-12* (called *SUT2*) (Hackel et al., 2006) and wild-type Money Maker plants (WT) were received from the Biology Department in Humboldt University of Berlin (Germany). AS lines showed a slower growth at the beginning of their development compared to WT, and sowing of AS plants was done 2.5 weeks earlier than WT, to obtain at grafting time a same height of the plants and number of expanded leaves (7 fully expanded). Eight and ten and a half weeks after sowing, respectively for WT and *SUT1*-AS and *SUT2*-AS lines, inoculation was performed by chip-grafting (Ahmad et al., 2014) with the strain PO of stolbur phytoplasma (STOL-PO), a strain isolated from **<u>P</u>**yrénées **<u>O</u>**rientales in South France (Ahmad et al., 2014) and belonging to ‘*Candidatus* Phytoplasma solani’ species (Quaglino et al., 2013) and with a scion from infected WT tomato plants. Non-infected controls were grafted with non-infected scions. For each genotype 8 plants were grafted with respectively infected and non-infected material for a total of 48 plants. In these conditions, at 17 days after chip-grafting (DAG), six new leaves have emerged on the grafted plants, irrespective of the genotype.

The severity of phytoplasma symptoms was recorded by a notation scale from 0 to 4, with class 0: no symptoms, class 1: beginning of curling of the leaflets, class 2: mild curling of leaflets, class 3: light yellowing of interveinal tissue of the leaflets and leaflet deformation, class 3.5: interveinal yellowing and severe curling and class 4: severe reduction of leaflet’s area, with leaflet chlorosis and a crook-shape, a characteristic of Stolbur disease on tomato (Ahmad et al., 2013).

### Identification of tomato gene sequences and design of primers

Tomato sequences were retrieved from NCBI (http://www.ncbi.nlm.nih.gov/) and Phytozome (https://phytozome.jgi.doe.gov/pz/portal.html) databases (Goodstein et al., 2012) and expression pattern was analyzed on the Bio-Analytic Resource (Toufighi et al., 2005) (http://bar.utoronto.ca/efp_tomato/cgi-bin/efpWeb.cgi). Information about selected genes is reported in Table S1 and primers used in real-time qPCR experiments are listed in Table S2. The efficiency of each primer couple was evaluated as described in Pfaffl (2001).

### Total RNA extraction and plant gene expression

For RNA extraction and for all analyses, plant samples used in the experiments were collected between 11:00 h and 12:00 h, two hours before the middle of the day. The first visible symptoms of infection appeared at 17 days after chip-grafting (DAG). At this stage, about six new leaves had emerged since grafting (L1 to L6), both on WT and *Le-SUT1* and *Le-SUT2* antisense lines (Fig. **1**) and the plants had reached the same height. Sampling was done at 18 DAG. Leaf material was sampled from leaves L1, L4 and L6 (Fig. **1**). Four plants were sampled per genotype, condition and leaf level. For RNA extraction from a leaf, the midrib of the leaflets 1, 2 and 3 were collected and frozen in liquid nitrogen. Total RNA was isolated following a TRIzol-based extraction (Invitrogen) and a DNase treatment (DNase I RNase-free, Thermo Fisher Scientific). The first strand cDNAs was synthesized with M-MLV reverse transcriptase starting from 1 µg of total RNA. Real-time qPCRs were performed on a CFX96 Real Time PCR Detection System (Bio-Rad Laboratories) using Takyon ROX SYBR 2X MasterMix dTTP blue (Eurogentec) beginning with a step at 95°C for 3 min, followed by 40 cycles for 15 s at 95°C, 60 s at 60°C and 30 s at 72°C. The mean normalized expression (MNE) was calculated by the method of normalization described in (Muller et al., 2002) using UBI, UPL3, PGK and UrK as reference genes and taking into account each primer couple efficiency (Table **S2**). Normalized data are expressed in relative units.

### Phytoplasma detection

Genomic DNA was extracted from 0.5 g of petiole and rachis from L1 leaf (Ll1 terminal leaflet), using Cetyl Trimethyl Ammonium Bromide method, 4 plants for each genotype and condition(Murray and Thompson, 1980). 200 ng of total leaf DNA were analyzed by real-time qPCR using primers for the *Methionine aminopeptidase* (*Map*) gene (Pelletier et al., 2009) (Table **S2**). Real-time qPCR was performed on a Light Cycler 480 (Roche) using SYBR^®^ Green master mix (Roche), imposing a step at 95°C for 15 min, followed by 45 cycles for 15 s at 94°C, 30 s at 62°C and 30 s at 66°C, then melting at 95°-10 s, 66°-10 s, continuous up to 95°. Absolute quantification was obtained with DNA fragment from the *Map* gene cloned in pGEM^®^-T Easy vector (Promega) with 10 to 10^8^ copies of the plasmid added on each PCR plate. rRNA abundance was analyzed to provide an additional index of phytoplasma multiplication. Using the same plants (4 plants for each genotype per condition), one ng of total RNA obtained from midribs of L1 leaf (leaflets 1 to 3) was analyzed by real-time qPCR using specific 16S rRNA primers and expressed in relative units (Table **S2**).

### Light microscopy

Seventy µm-thick transversal sections of the midrib of leaflets 4 or 5 from L1 leaf were cut using a vibratome (Leica) and stained with periodic acid (1% w/v, Sigma Aldrich) and Schiff’s reagent (VWR). Observations were carried out with Axiozoom V16 macroscope (Zeiss) equipped with a Plan-Neofluar Z 2.3x/0.57 RWD 10.6 mm objective. At least eight sections from four plants per genotype and condition were observed.

### Ultrastructure analysis using transmission electron microscopy

The midrib of leaflets 4 or 5 of L1 leaf was examined by transmission electron microscopy (TEM). Samples were fixed in 2.5% glutaraldehyde-2% paraformaldehyde in 100 mM phosphate buffer, pH 7.2, for 3 hours. They were post-fixed overnight at 4°C with 1% osmium tetroxide, then dehydrated in a graded ethanol series and progressively infiltrated with Epon resin for 48 h. Curing occurred 24 h at 60°C. 100 nm-thick sections were cut with an Ultracut S Microtome (Leica) and collected on hexagonal 600 mesh copper grids. Sections were observed at 120KV on a FEI Tecnai G2 Spirit TEM equipped with an Eagle 4K digital camera. Two plants per genotype were observed for infected plants and one for non-infected. For infected samples both transversal and longitudinal sections were analyzed. The number of peroxisomes in the phloem cells was counted per mesh (i.e. region of interest, ROI), one mesh corresponding approximately to 1000 µm^2^. Six to fourteen ROIs, focused on the phloem cells, including phloem parenchyma, phloem perivascular cells and companion cells, were observed per genotype and condition.

### Sugars and starch

Glucose, fructose and sucrose were quantified for the same leaflets 1, 2 and 3 of leaves L1, L4 and L6 of the four plants used for gene expression, using the leaf laminar tissues after removal of the midribs used for RNA extraction. Sugar quantification was assayed enzymatically (Enzytec™ Sucrose/D-Glucose/D-Fructose-R-Biopharm AG kit) (Vilaine et al., 2013). Starch quantification was determined after the release of Glucose by incubation with a-amylase and amyloglucosidase (Sigma Aldrich) (Vilaine et al., 2013). Four replicates were analyzed per leaf level, for each plant with 4 plants for each genotype per condition. Leaf samples were collected between 11:00 h and 12:00 h.

### Collection of phloem-enriched exudates

The apical leaflet from L3 was used for phloem EDTA-facilitated exudation (Beneteau et al., 2010). Immediately after cutting, the rachis of leaflet was re-cut in 10 mM HEPES adjusted to pH 7.5 with NaOH, 10 mM EDTA, pH 7.5 where it remained for 3-5 min. Then the rachis was immersed in 400 µl of the same buffer and placed in a dark box with high humidity for 4 hours for the exudation (from midday to 16:00 h). Tissue fresh weights (FW) were recorded at the end of the experiment to express the exudation rate per mg of FW. The exudates of 7 to 8 plants were analyzed for each genotype per infection condition.

### Analysis of phloem-enriched exudates

Amino acids in exudates were analyzed in an UPLC-PDA system as described in (Renault et al., 2010). The quantification of sugars, sugar alcohols and organic acids was carried out using a GC-FID device (Lugan et al., 2009). To normalize the data, we determined a content for each metabolite within all metabolites in a sample (Fig. S1), using the method originally developed for normalizing phloem exudates (Yesbergenova-Cuny et al., 2016). First, data were log2- transformed and then the value for each quantified metabolite in the profile was corrected with respect to the mean log-content of all metabolites for the replicate, with a locally weighted scatterplot smoothing (LOWESS), all using R software (http://www.r-project.org). This normalization is required for the identification of the metabolites whose proportion is modified in response to the infection. ANOVA tests were performed on this dataset after removing any metabolite with missing values. A given metabolite’s content was declared different when the adjusted *p*-value after a Benjamini-Hochberg correction was lower than 0.05.

### Statistical analysis

Statistical analyses on data sets including ANOVA were done using R statistical software. Correlations were calculated using the Pearson correlation coefficient and tested with the pairwise two-sided *p*-values and adjusted *p*-value determined with the Holm’s method. Hierarchical cluster analysis (HCA) graphical representation were done with Genesis version

1.7.6 (http://genome.tugraz.at/) after log2 transformation and normalization by the median, using complete linkage clustering option and Euclidean distance.

## Supplemental Material

Supplemental Figure S1: Workflow of metabolite data treatments.

Supplemental Figure S2: Transversal sections of apical leaves from non-infected and infected plants.

Supplemental Figure S3: Frequency and location of phytoplasmas in the sieve elements of infected plants.

Supplemental Figure S4: Details of the ultrastructure of the leaf main vein and peroxisome location in non-infected and infected plants.

Supplemental Figure S5: Amounts of sugars and starch in the leaves in response to the phytoplasma infection.

Supplemental Figure S6: Heat map of metabolite profiles of the phloem-enriched exudate from non-infected and infected plants in the three genotypes.

Supplemental Table S1: List and characteristics of candidate genes.

Supplemental Table S2: List of primers.

Supplemental Table S3: Severity of the symptoms due to ‘*Ca*. Phytoplasma solani’ infection in WT, *SUT1*-AS and *SUT2*-AS plants.

Supplemental Table S4: Exudation rate of metabolites identified in the phloem sap exudates of non-infected and infected plants.

Supplemental Table S5: Content of metabolites in the phloem sap exudates of non-infected and infected plants.

Supplemental Table S6: Two-way ANOVA of the expression of candidate genes in non-infected and infected plants.

Supplemental Table S7: Correlations in gene expression.

## Acknowledgments

We thank Christina Kühn for the gift of *LeSUT1*-AS and *LeSUT2*-AS seeds. FDM received the support of the EU Marie-Curie FP7 COFUND People Program, through an award of the AgreenSkills fellowship (under grant agreement n° 267196). Preliminary studies realized in this work benefited from the support of the BAP department of INRA (Vasculodrome’s project). We thank Catherine Jonard, Solenne Berardocco, Sylvain Déchaumet and Nathalie Marnet for the metabolite profiling at P2M2-IGEPP (INRA, Rennes). The IJPB benefits from the support of the LabEx Saclay Plant Sciences-SPS (ANR-10-LABX-0040-SPS). We thank the Imaging and Cytology platform of the Plant Observatory (IJPB, INRA Versailles-Grignon, France). We thank Priscilla Montfalet for initial development of the normalization method of the phloem sap exudates, Françoise Gilard for initial analysis of phloem-enriched exudates and Michael Hodges and Xavier Foissac for stimulating discussions. Imaging was performed at Bordeaux Imaging Center, a France BioImaging national infrastructure (ANR-10-INBS-04).

## List of Authors contributions

S.D., F.D.M and S.E. conceived and supervised the experiments. F.D.M. performed observations by light microscopy, gene expression analysis, soluble sugars quantification with the contribution of R.L.H and F.V. B.B. performed transmission electron microscopy observations. F.R. and S.E. performed phytoplasma detection analyses. S.D. analyzed metabolic profiles with contribution of A.B. and M.L M.M. Writing and editing of the manuscript was done by F.D.M., M.R.T. and S.D. All authors contributed to the corrections. S.D. agrees to serve as the author responsible for contact and ensures communication.

## Abbreviations

SE: sieve element, CC: companion cell, ICL: isocitrate lyase, MLS: malate synthase, GLO: glycolate oxidase, FRK: fructokinase, GLYR; glycolate reductase. WT: Wild- type, AS: antisense, TCA: tricarboxylic acid cycle. SUT1: sucrose transporter 1, SUT2: sucrose transporter 2.

## Legend of the supplementary Figures

**Supplemental Figure S1.**
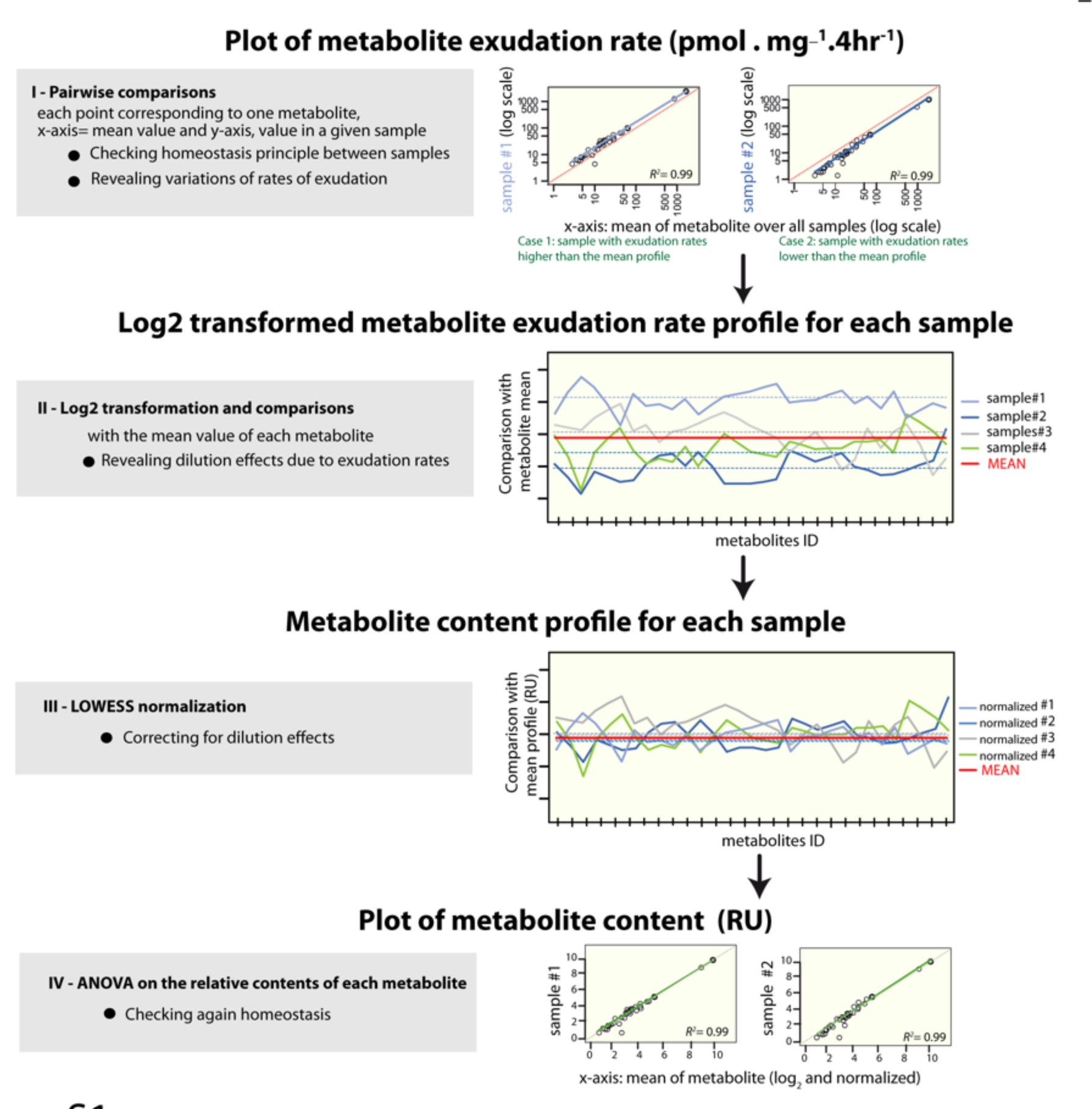
Workflow of metabolite data treatments. Workflow of phloem-enriched exudate metabolite analysis, consisting of data pairwise comparisons to verify homeostasis between samples, log2 transformation, LOWESS normalization and statistical analysis. This procedure reveals the variations of exudation rates and corrects for dilution effects. RU: relative units

**Supplemental Figure S2.**
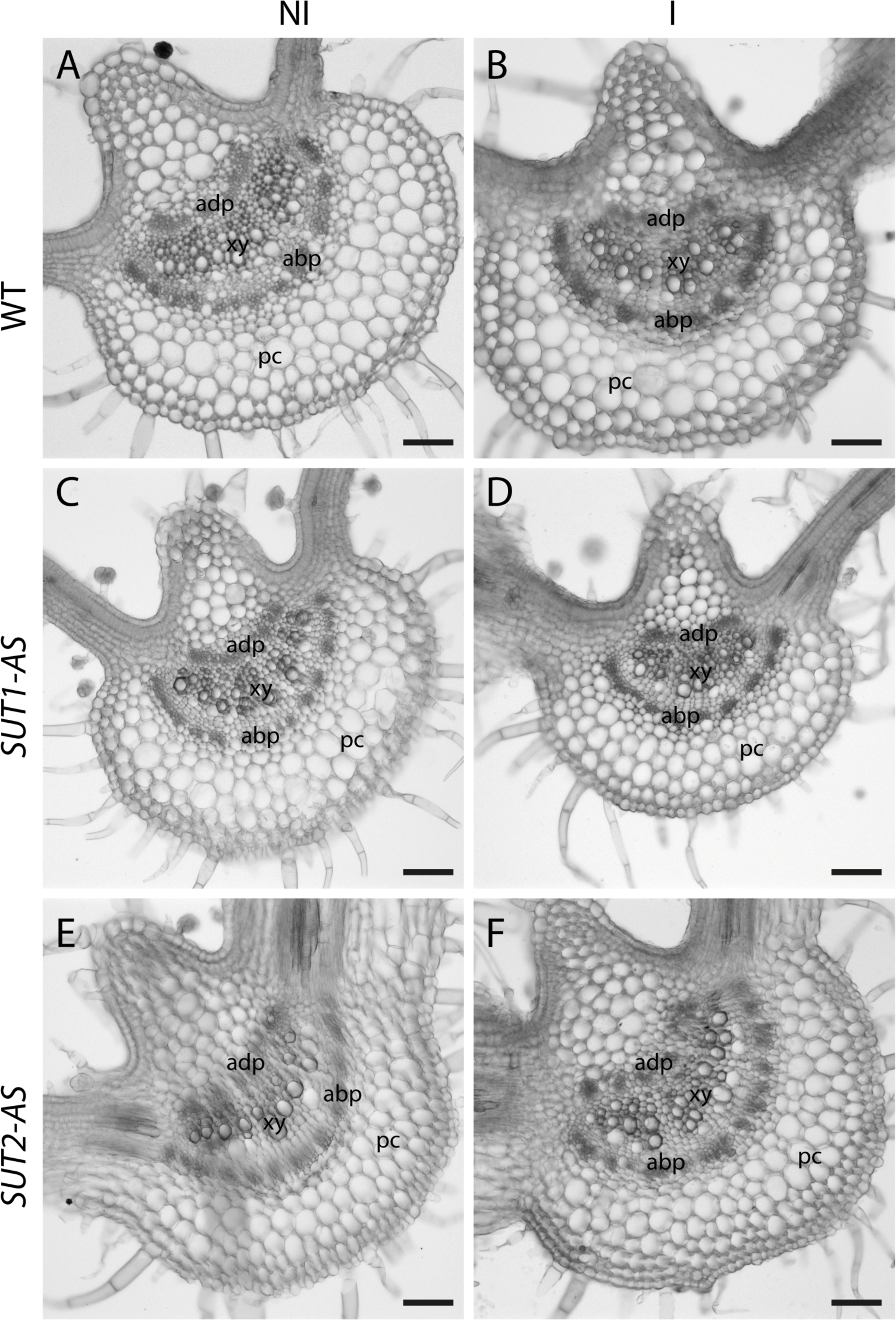
Transversal sections of L1 leaves from not-infected and infected plants. Images were obtained on fresh transverse thin sections of the main vein of L1 leaflet. A,B: Wild-type (WT), C,D: *SUT1*-AS, E,F: *SUT2*-AS transversal main vein sections of not-infected (NI) and infected (IF) plants respectively. Vascular tissues exhibit a typical organization with xylem vessels surrounded by abaxial and adaxial phloem. adp, adaxial phloem, abp, abaxial phloem, xy, xylem, pc, parenchyma cells. Bar, 100 µm.

**Supplemental Figure S3.**
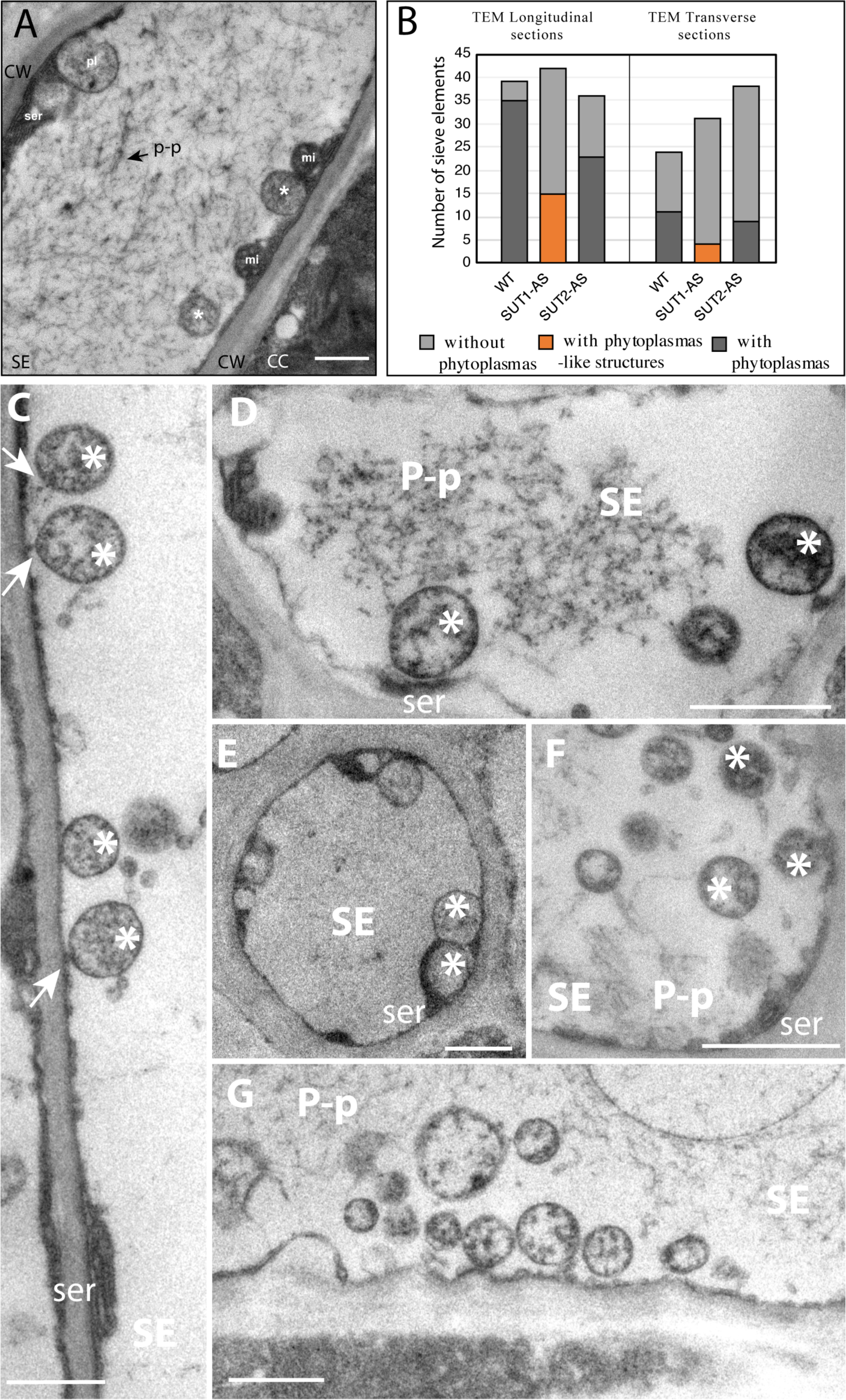
Frequency and location of phytoplasmas in the sieve elements of infected plants. **A, C to G**: Details of Transmission Electron Microscopy (TEM) micrographs in the sieve elements of infected main veins of L1 (leaflets 4 or 5) showing the location of parietal phytoplasma. **B:** Frequency of sieve elements with phytoplasmas. **A:** Distinctive features of phytoplasma observed with TEM. Observation of phytoplasmas, plastids and mitochondria on a sieve tube longitudinal section. A mature sieve-tube plastid (pl), around 1 µm wide, exhibits a sparse stroma enclosing a dense inclusion of proteinaceous type. Phytoplasma (asterisks), less wide, display a loose fibrillar content whereas mitochondria matrix (mi) is dense with clearer cristae. Bar: 1 µm. **B**: Number of sieve elements with or without phytoplasmas observed in the phloem of L1 leaves from infected plants. The data were determined with transmission electron microscopy images of transverse or longitudinal sections of the phloem of WT, *SUT1*-AS and *SUT2*- AS. A total of 63, 73 and 74 SEs were imaged for WT, *SUT1*-AS and *SUT2*-AS plants, respectively. Phytoplasmas were unambiguously identified in the SEs of WT and *SUT2*-AS plants (5-30 phytoplasmas per cell) while they were difficult to identify in *SUT1*-AS plants, where they look like vesicular structures (1-2 phytoplasma-like structure per cell). More phytoplasma were observed on longitudinal sections than on transversal sections, likely due to an unequal distribution in the sieve elements, (see for example Fig. 3I), with frequent accumulation of phytoplasma at one side of sieve plates. **C,D**: TEM images of Wild-type (WT), **E,F**: TEM images of SUT1 antisense line (AS), **G**: TEM images of *SUT2*-AS antisense line. White arrows indicate attachments of phytoplasma bodies to the sieve-element plasma membrane. *: Phytoplasma, SE, sieve element, P-p: filamentous P-proteins, ser: sieve element reticulum. Bar: 1 µm.

**Supplemental Figure S4.**
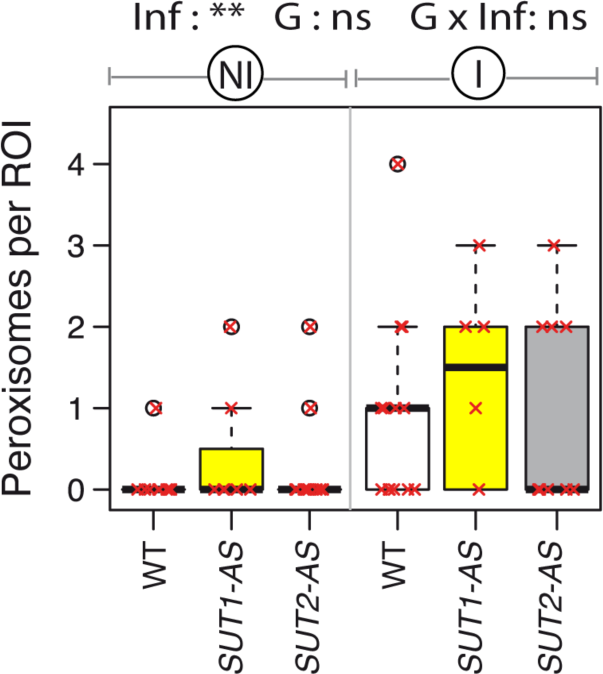
Frequency of peroxisomes in the phloem of non-infected and infected tomato plants. Frequency of peroxisomes determined on TEM images obtained from not-infected plant and infected plants. Above the boxplot, significance of the effects due to the infection (Inf), genotype (G), and their interaction (G x Inf), determined using a two-way ANOVA (*, *P* < 0.05; **, *P* < 0.01; ***, *P* < 0.001; ns, not significant). Stars above the boxes indicate significant differences by a *t*-test in *SUT1*- or *SUT2*- AS plants compared to WT plants in the same conditions (*, *P* < 0.05; **, *P* < 0.01; ***, *P* < 0.001; ns, not significant). ROI: region of interest. White boxes: Wild-type (WT); light grey: *SUT1*-AS line (AS); dark grey: *SUT2-*AS. The boxplots show the number of peroxisomes per ROI with *n*=6 to 14 depending on conditions. The red crosses indicate the value for each replicate.

**Supplemental Figure S5.**
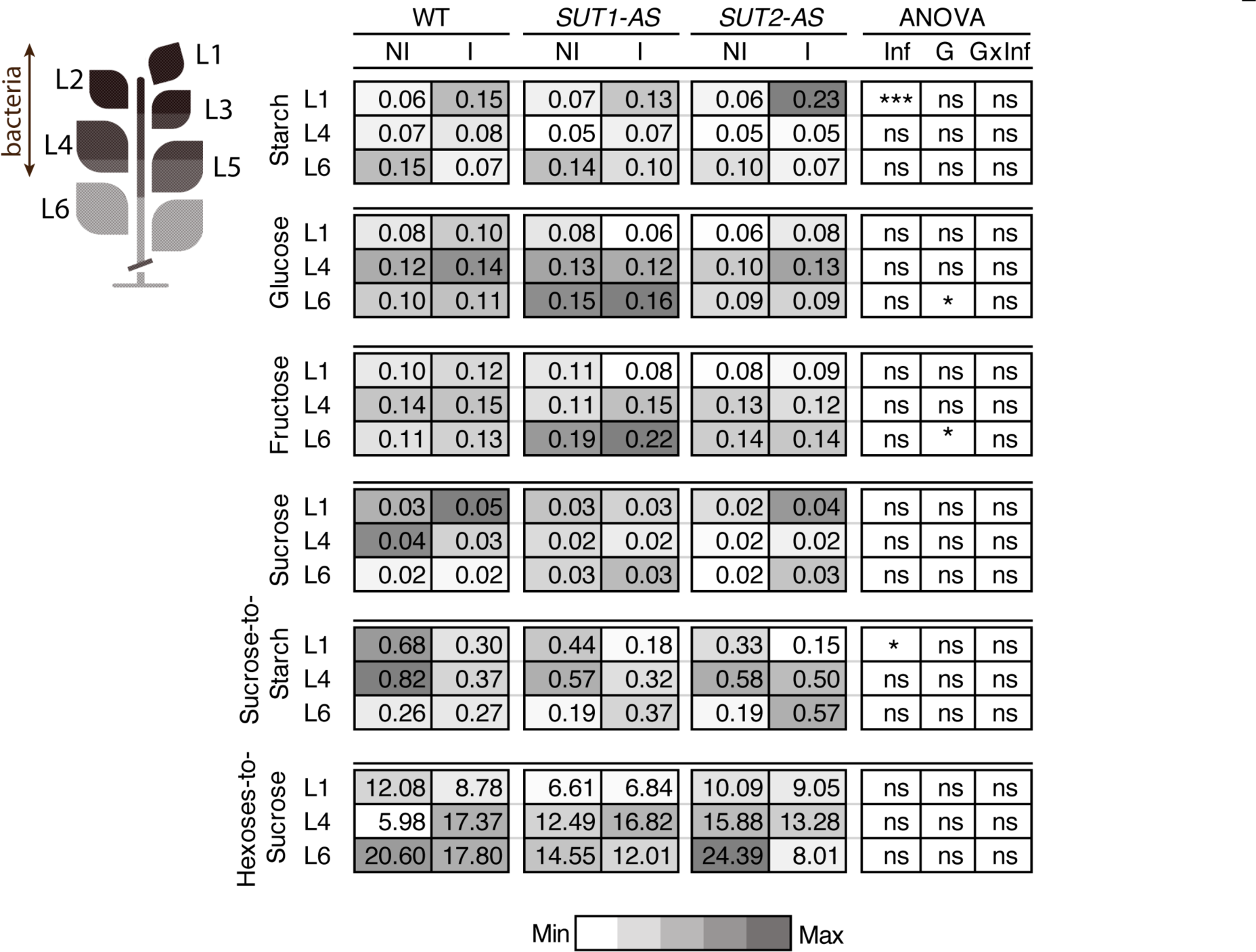
Amounts of sugars and starch in the leaves in response to the phytoplasma infection. Heat map and ANOVA for starch and soluble sugar contents at different leaf levels. The data were obtained on L1, L4 and L6 leaves sampled at 18 days after grafting. Left panel: the mean values for each compound and each genotype in not-infected (NI) and infected (I) plants. Data are expressed in nmol mg^-1^ of sample fresh weight. Right panel: *p*-values obtained by a two-way ANOVA for each leaf level (*, *P* < 0.05; **, *P* < 0.01; ***, *P* < 0.001; ns, not significant), with Inf: infection effect, G: genotype effect and G x Inf: genotype per infection interaction effect.

**Supplemental Figure S6.**
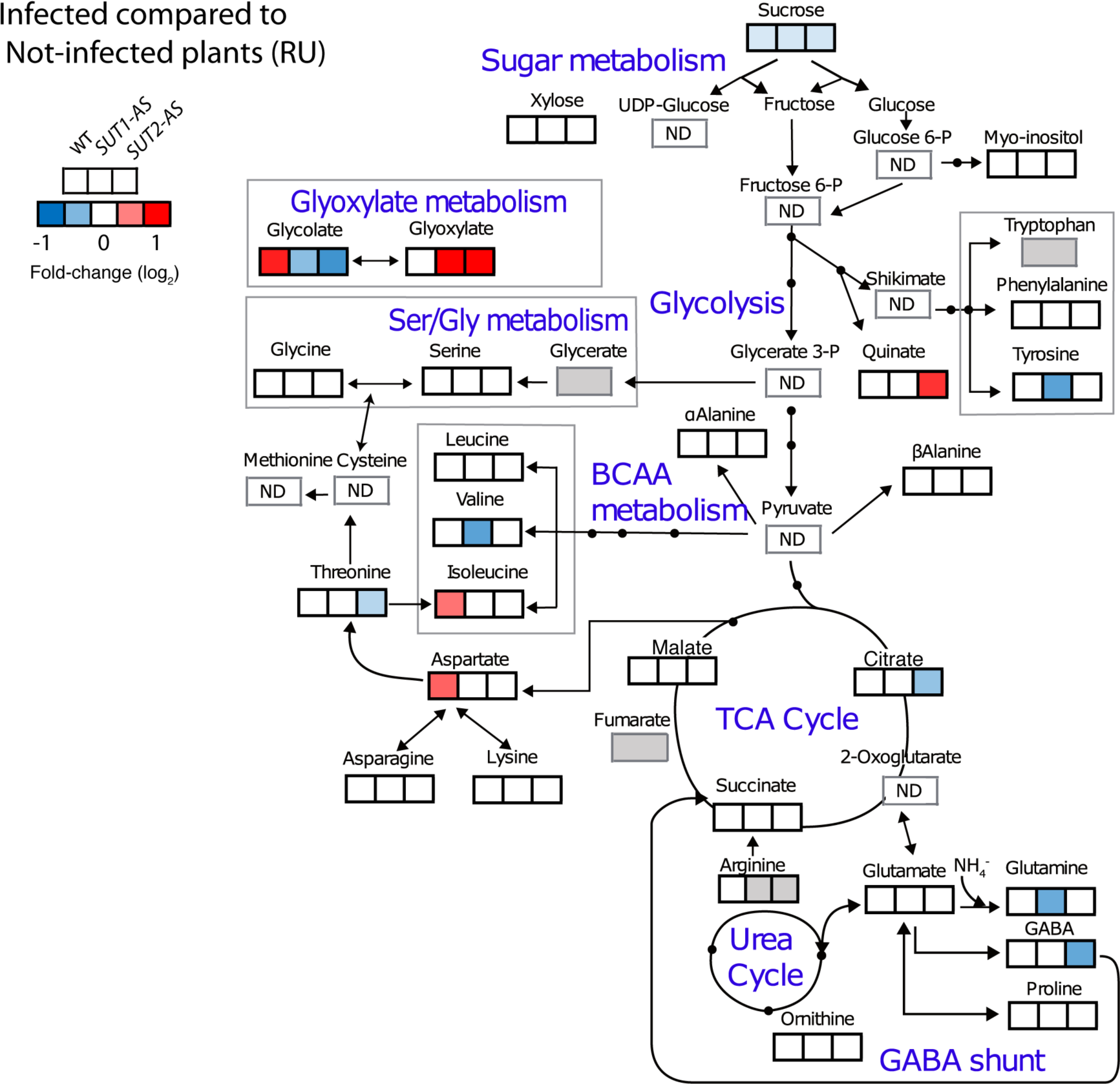
Comparison of metabolite profiles of the phloem-enriched exudate from the L3 leaf of not-infected and infected plants in the three genotypes. Fold changes in the content of metabolites in phloem-enriched exudates in response to infection. Inserted heat maps represent significant fold-changes (*P*value < 0.05 on a paired t-test) between infected and not-infected plants. From the left to the right sides, responses in wild-type (WT), *SUT1*-AS and *SUT2*- AS plants. Values are shown in a blue-to-red log2 scale with blue values for metabolites showing a smaller content (and red values for higher) due to infection. BCAA: branched-chain amino acids. TCA: tricarboxylic acid cycle. In white: non-significant variations. In grey: missing values.

**Table S1.** List and characteristics of candidate genes.

| Gene name | Arabidopsis ortholog | Function | Expression* | References |
| --- | --- | --- | --- | --- |
| Defence related genes |  |  |  |  |
| *PR1a/P4* | At4g33720 (putative *PR1*) | Pathogenesis related protein | l<br>SA dependent defence gene, upregulation by *Ca. P. s.* infection | van Kan *et al.* (1992); Ahmad *et al.* (2014); Ahammed *et al.* (2018) |
| *PR2a* | At4g16260 (β-1,3-glucanase) | Pathogenesis related protein (β-1,3-glucanase) | l<br>SA dependent defence gene, upregulation by *Ca. P. s.* infection | (van Kan *et al.* (1992); Ahmad *et al.*, 2014) |
| *CAS2* | At2g31960 (GLS03) | Putative callose synthase | l<br>No response to *Ca. P. s.* infection | De Marco, personal communication |
| *CAS7* | At1g06490 (*CAS7*) | Putative callose synthase | l<br>Upregulation by *Ca. P. s.* infection | (De Marco *et al.*, 2016) |
| Vascular marker |  |  |  |  |
| *PP2* | At4g19840 (*PP2-A1*) | Phloem protein 2 (lectin) | l, phloem<br>Downregulation by *Ca. P. s.* infection | (De Marco *et al.*, 2016) |
| Sugar metabolism |  |  |  |  |
| *FRK1* | At5g51830 (*FRK1*) | Cytosolic fructokinase | l, st, r, f, vascular tissues | Kanayama *et al.* (1997); Damari-Weissler *et al.* (2006); Stein *et al.* (2018) |
| *FRK2* | At1g06030 (*FRK2/6*) | Cytosolic fructokinase | s, r, f, vascular tissues<br>Increased protein amount in response to *R.s.* | Damari-Weissler *et al.* (2006, 2009); Dahal *et al.* (2009) |
| *FRK3* | At1g66430 (*FRK3/6*) | Plastidial fructokinase | r, l, st, s. vascular tissues.<br>Increased protein amount in response to TCMV infection | German *et al.* (2004); Damari-Weissler *et al.* (2006); Stein *et al.* (2016); Carmo *et al.* (2017) |
| *SUSY1* | At5g20830 (*SUS1*) | Sucrose synthase | st, vascular tissues (xylem) | Goren *et al.* (2011) |
| *SUSY3* | At3g43190 (*SUS4*) | Sucrose synthase | r | Goren *et al.* (2011) |
| Sugar transport |  |  |  |  |
| *SUT1* | At1g22710 (*SUC2*) | Sucrose transporter | l, st, source organs<br>mRNA in CC, protein in SE | Kuhn *et al.* (1996); Hackel *et al.* (2006); Zhao *et al.* (2018) |
| *SUT2* | At2g02860 (*SUC3*) | Sucrose transporter | l, st, sink organs<br>mRNA in CC, Protein in SE. Upregulation by *RKN* | (Barker *et al.* (2000); Zhao *et al.* (2018) |
| *SWEET2a* | At3g14770 (*SWEET2*) | Sugar facilitator | l, st, r, fl<br>Upregulation by *RKN* | Feng *et al.* (2015); Zhao *et al.* (2018) |
| *SWEET5b* | At5g62850<br>(*SWEET5*) | Sugar facilitator | l, fl<br>Upregulation in leaves<br>and roots by *RKN* | Feng *et al.* (2015);<br>Zhao *et al.* (2018) |
| *SWEET10c* | At5g50790<br>(*SWEET10*) | Sugar facilitator | l, fl<br>Upregulation by sugars,<br>salt and temperature | Feng *et al.* (2015) |
| *SWEET11a* | At3g48740<br>(*SWEET11*) | Sugar facilitator | l, fl<br>Upregulation by sugars,<br>salt and temperature | Feng *et al.* (2015) |
| *SWEET12a* | At5g23660<br>(*SWEET12*)<br>At3g48740<br>(*SWEET11*) | Sugar facilitator | l, fl<br>Upregulation by sugars,<br>salt and temperature | Feng *et al.* (2015) |
| Glyoxylate cycle and photorespiration |  |  |  |  |
| *ICL* | At3g21720<br>(*ICL*) | Isocitrate lyase | yl, sl. Protein in yl. and<br>sl., less in mat. leaves | Nieri *et al.* (1997);<br>Famiani *et al.* (2016) |
| *MLS* | At5g03860<br>(*MLS*) | Malate synthase | yl, sl. Protein in yl. and<br>sl., lower activity in the<br>roots by *R.s.* infection | Nieri *et al.* (1997);<br>Famiani *et al.* (2016);<br>Fan *et al.* (2018) |
| *GLO40/GOX1/*<br>*SIGlo1/TomGlo.1* | At3g14420<br>(*GOX1*),<br>At3g14415<br>(*GOX2*) | Glycolate oxidase | l<br>Decreased activity in<br>response to *F. l.*. No<br>response to *P.s.* | Sanwal & Waygood,<br>(1963); Speirs *et al.*<br>(1995); Ahammed *et al.* (2018) |
| *GLO00/GOX2*<br>*/SIGlo2* | At3g14420<br>(*GOX1*)<br>At3g14415<br>(*GOX2*) | Glycolate oxidase | yf, l<br>Decreased activity in<br>response to *F.s.*<br>Upregulation by *P.s.*<br>Involved in the<br>photorespiration | Sanwal & Waygood<br>(1963); Ohta *et al.*<br>(2005); Ahammed *et al.* (2018) |
| *GLO50/GOX3/SIGlo3* | At4g18360<br>(*GOX3*) | Glycolate oxidase | l<br>Decreased activity in<br>response to *F. l.* | Sanwal & Waygood<br>(1963) |
| *GLYR1* | At3g25530<br>(*GLYR1*) | Glyoxylate<br>reductase | l<br>Increased activity by *F. l.*<br>infection | Sanwal & Waygood<br>(1963) |
| *GLYR2* | At1g17650<br>(*GLYR2*) | Glyoxylate<br>reductase | l<br>Increased activity by *F. l.*<br>infection | Sanwal & Waygood<br>(1963) |

\* Gene pattern of expression, unless specifically indicated, with: l: leaf, yl: young leaf, sl: senescent leaf, s: stem, r: roots, s: seeds, yf: young fruit, mf: mature fruits, f: fruit, fl: flowers. TCMV: tomato chlorotic mottle virus infection. *P.s.*: *Pseudomonas syringae* pv. tomato DC3000, *F.s.*: *Fusarium lycopersici*. *R. s.*: *Ralstonia solanacearum*. *Ca. P. s.*: 'Candidatus Phytoplasma solani', *RKN*: root-knot nematode. *CC*: companion cells. *SE*: sieve elements.

**Table S2.**
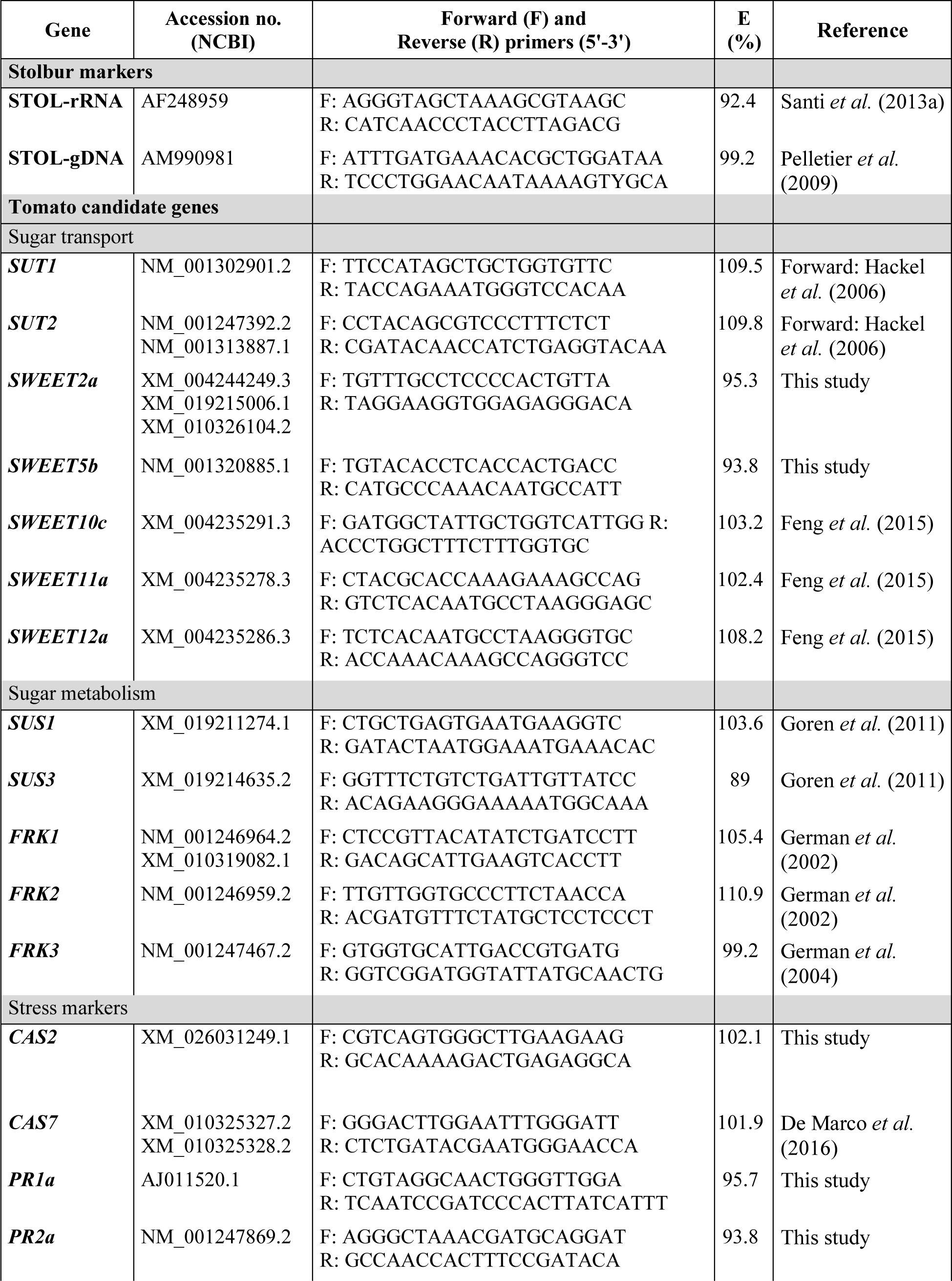
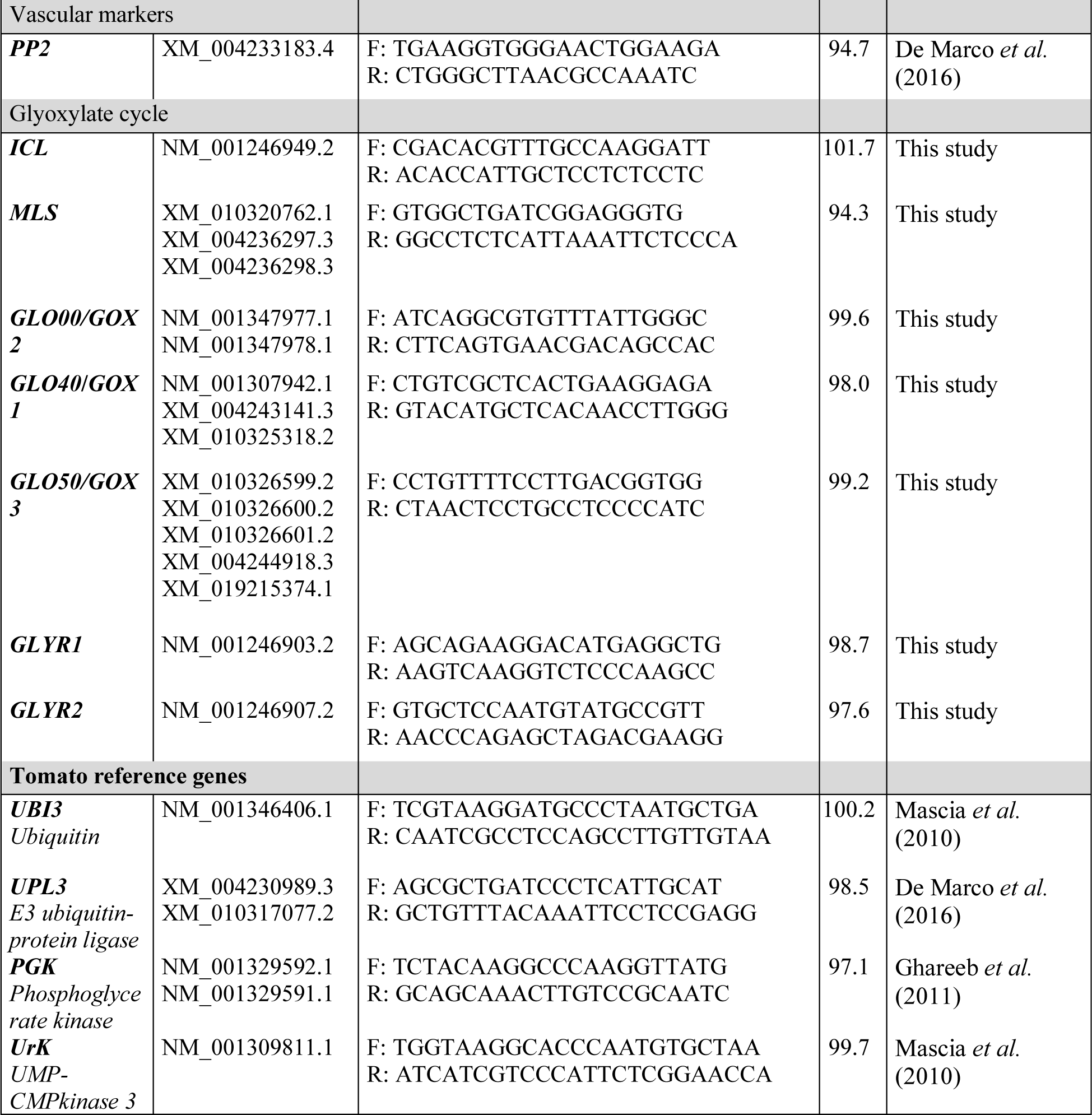
List of Primers. When more than one sequence was identified, primers amplify each variant. E (%): Efficiency of the primers.

**Table S3.** Symptoms after infection by ‘*Ca*. Phytoplasma solani’ in WT, *SUT1*-AS and *SUT2*-AS plants. Number of plants for each class of symptoms. The classes of symptoms, from 0 to 4, correspond to no symptoms (class 0) to small crook-shaped and chlorotic leaves (class 4) (see details in the main text). Data were obtained on 4 plants at 3 dates after grafting (DAG: days after grafting). Above panel: number of plants for each class of symptoms. Bottom panel: Kruskal Wallis tests followed by Dunn test performed on the symptoms level (classes from 0 to 4) for each date.

|  | WT | *SUT1*-<br>AS | *SUT2*-<br>AS | WT | *SUT1*-<br>AS | *SUT2*-<br>AS | WT | *SUT1*-<br>AS | *SUT2*-<br>AS |
| --- | --- | --- | --- | --- | --- | --- | --- | --- | --- |
| Class | 18 DAG |  |  | 24 DAG |  |  | 27 DAG |  |  |
| 0 | 0 | 3 | 0 | 0 | 0 | 0 | 0 | 0 | 0 |
| 1 | 0 | 1 | 0 | 0 | 0 | 0 | 0 | 0 | 0 |
| 2 | 1 | 0 | 0 | 0 | 2 | 0 | 0 | 1 | 0 |
| 3 | 1 | 0 | 4 | 1 | 2 | 2 | 0 | 2 | 1 |
| 3.5 | 0 | 0 | 0 | 0 | 0 | 0 | 1 | 1 | 0 |
| 4 | 2 | 0 | 0 | 3 | 0 | 2 | 3 | 0 | 3 |

| Kruskal Wallis test *p*-value | 18 DAG | 24 DAG | 27 DAG |
| --- | --- | --- | --- |
|  | 0.0207 * | 0.0505 | 0.0455 * |

| Post hoc Dunn test adjusted with the Holm method | 18 DAG | 24 DAG | 27 DAG |
| --- | --- | --- | --- |
| WT vs *SUT1*-AS | 0.0304 * | 0.0592 | 0.069 |
| WT vs *SUT2*-AS | 0.7188 | 0.5963 | 0.7915 |
| *SUT1*-AS vs *SUT2*-AS | 0.054 | 0.1433 | 0.0891 |

**Table S4.**
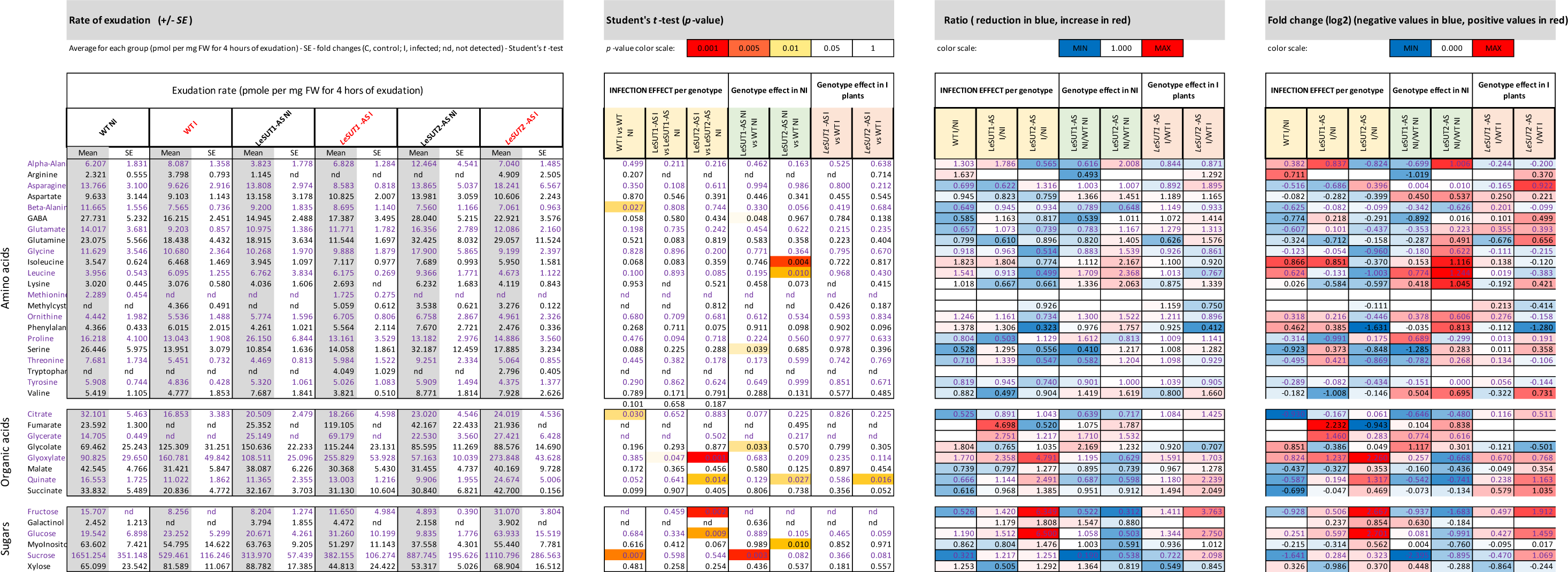
Rate of exudation for each metabolites detected in the phloem exudates.

**Table S5.**
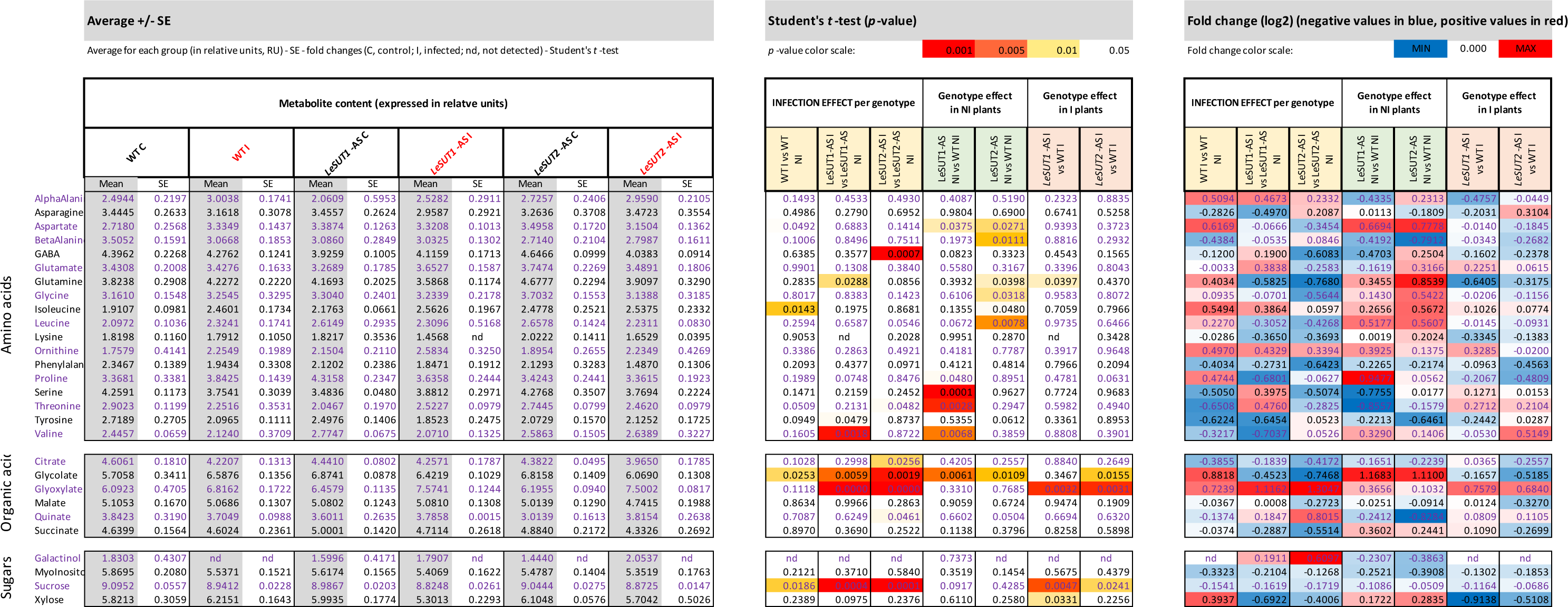
Relative content of the metabolites identified in the phloem sap exudates (log2 transformed and normalized values).

**Table S6.** Two-way ANOVA of the expression of candidate genes in not-infected and infected plants. The ANOVA was determined on gene expression data obtained at 3 leaf levels for the 3 genotypes (WT, *SUT1*-AS, *SUT2-*AS). G: Genotype; L: leaf position; GxL= Genotype per leaf position interaction. *p*-value (2-way ANOVA): *, *P* <0.05, **; *P* <0.01; ***, *P* <0.001. *n*=72

| Gene | G | L | GxL | G | L | GxL |
| --- | --- | --- | --- | --- | --- | --- |
|  | Not-infected |  |  | Infected |  |  |
| *SUS1* | 0.0132 * | 4.07e-05 *** | 0.4750 | 0.2085 | 3.83e-05 *** | 0.6739 |
| *SUS3* | 0.2073 | 0.0053 ** | 0.7242 | 0.0057 ** | 0.0006 *** | 0.0888 |
| *FRK1* | 0.1837 | 0.0004 *** | 0.6185 | 0.0004 *** | 8.59e-05 *** | 0.6609 |
| *FRK2* | 0.0017 ** | 0.0085 ** | 0.2882 | 0.3586 | 0.0001 *** | 0.5379 |
| *FRK3* | 0.0226 * | 0.0031 ** | 0.3118 | 0.1911 | 0.0033 ** | 0.5541 |
| *CAS2* | 0.0055 ** | 2.16e-07 *** | 0.0239 * | 0.0124 * | 2.63e-06 *** | 0.3743 |
| *CAS7* | 0.0747 | 2.32e-10 *** | 0.9559 | 0.5643 | 4.46e-06 *** | 0.6072 |
| *PR1a* | 0.0584 | 0.0854 | 0.5372 | 0.0122 * | 0.2525 | 0.3696 |
| *PR2a* | 0.4755 | 0.0265 * | 0.4345 | 0.1518 | 0.2261 | 0.5719 |
| *SWEET2a* | 0.0089 ** | 0.0008 *** | 0.5491 | 0.0268* | 0.0016** | 0.5703 |
| *SWEET5b* | 0.1581 | 6.03e-06 *** | 0.7338 | 0.3010 | 9.37e-06 *** | 0.0906 |
| *SWEET10c* | 0.2010 | 0.0007 *** | 0.5131 | 0.4201 | 5.20e-05 *** | 0.8110 |
| *SWEET11a* | 0.4628 | 0.0033 ** | 0.4946 | 0.0417 * | 3.57e-05 *** | 0.7356 |
| *SWEET12a* | 0.3217 | 0.0004 *** | 0.5877 | 0.0497 * | 0.0001*** | 0.8447 |
| *PP2* | 0.5485 | 3.74e-07 *** | 0.5769 | 0.472 | 2.79e-06 *** | 0.6045 |
| *ICL* | 0.0640 | 0.0370 * | 0.6605 | 0.0145 * | 0.3298 | 0.5561 |
| *MLS* | 0.0568 | 0.4368 | 0.9470 | 0.0002 *** | 0.9635 | 0.9771 |
| *GLO00/GOX2* | 0.0035 ** | 0.3770 | 0.6526 | 0.0142 * | 0.0025 ** | 0.4767 |
| *GLO40/GOX1* | 0.5239 | 0.0206 * | 0.9951 | 0.3176 | 0.0009 *** | 0.7862 |
| *GLO50/GOX3* | 0.2170 | 0.2787 | 0.9831 | 0.5420 | 0.1109 | 0.5444 |
| *GLYR1* | 0.4658 | 0.0318 * | 0.4585 | 0.6818 | 0.0355 * | 0.6063 |
| *GLYR2* | 0.0012 ** | 0.0025 ** | 0.0951 | 0.0356 * | 0.6020 | 0.0756 |

**Table S7.** Correlations in gene expression. The *Pearson* correlation was determined on the full gene expression dataset obtained at the three leaf levels (L1, L4 and L6) of the infected and non-infected plants of the 3 genotypes (WT, *SUT1*-AS and *SUT2-*AS). *n*=72.

| Correlations | *Pearson*<br>correlation | Pairwise two-sided *p*-values | Adjusted<br>*p*-values |
| --- | --- | --- | --- |
| Glycolysis and peroxisome metabolism |  |  |  |
| *FRK1* - *ICL* | -0.406 | 0.000 | 0.019 |
| *FRK1* - *GLYR2* | -0.384 | 0.001 | 0.037 |
| *FRK3* - *GLO40/GOX1* | 0.394 | 0.009 | 0.028 |
| *FRK3* - *GLO50* | 0.416 | 0.005 | 0.014 |
| *FRK3* - *GLYR1* | 0.401 | 0.030 | 0.022 |
| *SUS1* - *GLO40/GOX1* | 0.438 | <0.0001 | 0.006 |
| *SUS3* - *GLO40/GOX1* | 0.450 | <0.0001 | 0.004 |
| *SUS3* - *GLO50/GOX3* | 0.470 | <0.0001 | 0.002 |
| *SUS3* - *GLYR1* | 0.473 | <0.0001 | 0.001 |
| Glycolysis |  |  |  |
| *SUS1* - *FRK1* | 0.446 | <0.0001 | 0.004 |
| *SUS1* - *FRK2* | 0.649 | <0.0001 | <0.0001 |
| *SUS1* - *FRK3* | 0.687 | <0.0001 | <0.0001 |
| *SUS1* - *SUS3* | 0.518 | <0.0001 | 0.000 |
| *SUS3* - *FRK2* | 0.486 | <0.0001 | 0.001 |
| *SUS3* - *FRK3* | 0.599 | <0.0001 | <0.0001 |
| *FRK1* - *FRK2* | 0.527 | 0.000 | 0.000 |
| *FRK2* - *FRK3* | 0.724 | 0.000 | 0.000 |
| Peroxisomal metabolism |  |  |  |
| *GLO00/GOX2* - *ICL* | 0.644 | <0.0001 | <0.0001 |
| *GLO00/GOX2* - *MLS* | 0.835 | <0.0001 | <0.0001 |
| *ICL* - *MLS* | 0.734 | <0.0001 | <0.0001 |
| *GLO50/GOX3* - *GLO40/GOX1* | 0.626 | <0.0001 | <0.0001 |
| *GLO50/GOX3* - *GLYR1* | 0.653 | <0.0001 | <0.0001 |
| *GLO40/GOX1* - *GLYR1* | 0.884 | <0.0001 | <0.0001 |
| Others |  |  |  |
| *CAS7* - *PP2* | 0.906 | <0.0001 | <0.0001 |
| *CAS7* - *FRK3* | 0.734 | <0.0001 | <0.0001 |
| *PP2* - *FRK3* | 0.742 | <0.0001 | <0.0001 |

